# Automated Modeling of Protein Accumulation at DNA Damage Sites using qFADD.py

**DOI:** 10.1101/2021.03.15.435501

**Authors:** Samuel Bowerman, Jyothi Mahadevan, Philip Benson, Johannes Rudolph, Karolin Luger

## Abstract

Cells are exposed to a plethora of influences that can cause damage to DNA and alter the genome, often with detrimental consequences for health. Cells mitigate this damage through a variety of repair protein pathways, and accurate measurement of the accumulation, action, and dissipation timescales of these repair proteins is required to fully understand the DNA damage response. Recently, we described the Q-FADD (<u>Q</u>uantitation of <u>F</u>luorescence <u>A</u>ccumulation after <u>D</u>NA <u>D</u>amage) method, which enhances the analytical power of the widely used laser microirradiation technique. In that study, Q-FADD and its preprocessing operations required licensed software and a significant amount of user overhead to find the model of best fit. Here, we present “qFADD.py”, an open-source implementation of the Q-FADD algorithm that is available as both a stand-alone software package and on a publicly accessible webserver (https://qfadd.colorado.edu/). Furthermore, we describe significant improvements to the fitting and preprocessing methods that include corrections for nuclear drift and an automated grid-search for the model of best fit. To improve statistical rigor, the grid-search algorithm also includes automated simulation of replicates. As an example, we discuss the recruitment dynamics of the signaling protein PARP1 to DNA damage sites, and we show how to compare different populations of qFADD.py models.

**Statement of Significance:** Cells are constantly bombarded by factors that can alter or damage their genome, and they have evolved a variety of proteins that can identify and fix this damage. To fully understand how these proteins interact in repair pathways, we need robust methods to quantify the timescales between the initial identification of the DNA damage event and the subsequent protein-protein interactions that lead to repair. Laser microirradiation is a popular method for studying these repair protein cascades in vivo, and methods for quantifying the timescales of recruitment in these experiments have historically been simple to implement but lacking in physical interpretation. Here, we present qFADD.py, the next iteration of the Q-FADD method, which uses Monte Carlo diffusion models to interpret repair protein recruitment timescales to sites of DNA damage. By moving towards automated fitting procedures with minimal bias from the user, qFADD.py provides a statistically robust but low-effort means to analyze laser microirradiation experiments through a biophysical framework.

## 1. Introduction

Living cells are constantly bombarded by DNA damaging agents, from sources both within and outside the cell. This damage can take many forms, such as double-strand DNA breaks, single-strand breaks, or base oxidation events(1–3). If unchecked, the resulting base changes (mutations), deletions, or chromosome fusion and breakage events result in permanent changes in the genome, with often detrimental effects for the cell. To protect from these damage events, cells have evolved a complex array of DNA repair pathways that utilize a wide variety of signaling and repair enzymes, each with varying activities throughout the cell cycle(4). While specific pathways counteract particular modes of damage, some proteins can participate in several repair processes, and inhibiting one repair cascade can alter the typically conserved sequential accumulation of signaling and repair proteins(5).Improper regulation of DNA damage sensing and repair proteins is strongly correlated with, and can even result in, several cancers(6–8). To this end, a better understanding of DNA damage pathways and the dynamics of the participating proteins is important to develop better cancer therapeutics, many of which function by introducing excessive DNA damage in rapidly growing cancer cells. As such, suppressing DNA damage repair in these cells will lead to more effective cancer drugs.

Accurate measurement of accumulation kinetics for the many proteins involved with signaling and repairing DNA damage events is necessary to understand the complex interplay within and between the various repair pathways. Laser microirradiation is one popular in vivo technique to study these kinetics due to its ability to track protein dynamics as a direct response to locally-induced and varied damage(9). In this method, cells are transfected to express one or several fluorescently-labeled proteins (depending on microscope capability). A defined region of chromatin is damaged with a short-wavelength (∼400 nm) laser and then monitored for time-dependent accumulation of the labeled protein(s) via fluorescence microscopy(10–12). Historically, these data have been fit with such methods as: determining the time required for half of the total accumulation (t1/2), fitting the timeseries to a single exponential, and fitting the timeseries with multiple exponentials(5, 12, 13). While straightforward to implement, we have previously shown that these methods lack the ability to differentiate between different nuclear shape profiles and therefore suffer from averaging diverse nuclei in to a single, representative timeseries(11). Moreover, as they are not direct measures of diffusion, they cannot be directly compared to orthogonal diffusion methods such as FRAP or fluorescence correlation spectroscopy (FCS)(9, 14, 15).

In response, we recently developed the “Quantitation of Fluorescence Accumulation after DNA Damage” (Q-FADD) approach(11). Q-FADD models protein motion as a free-diffusion process and can therefore be compared to other diffusion-based methods. Furthermore, nuclear shape and size is explicitly represented in the fitting process, which was shown to be an improvement over the t1/2 and exponential fitting methods. Originally, Q-FADD was conducted through a combination of MatLab and Mathematica notebooks, but here we describe “qFADD.py” (Figure 1) - an open-source Python implementation of the Q-FADD workflow that exhibits several significant improvements and updates compared to the original method. The new companion program (“image_analyzer.py”) can now properly account for nuclear drifting before converting captured image stacks into qFADD.py-readable nuclear envelope, ROI mask, and accumulation timeseries files, thereby reducing the number of microirradiation datasets that are discarded from analysis. An additional improvement through the qFADD.py implementation is a grid-search algorithm that automatically generates and compares multiple simulated model replicates over a wide range parameter combinations, reducing the potential for user-bias and increasing the statistical certainty of reported best-fit models. Laser microirradiation data of the poly(ADP-ribose) polymerase 1 (PARP1) signaling protein are presented as examples for the pipeline, and the interpretation of qFADD.py outputs are discussed.

**Figure 1.**
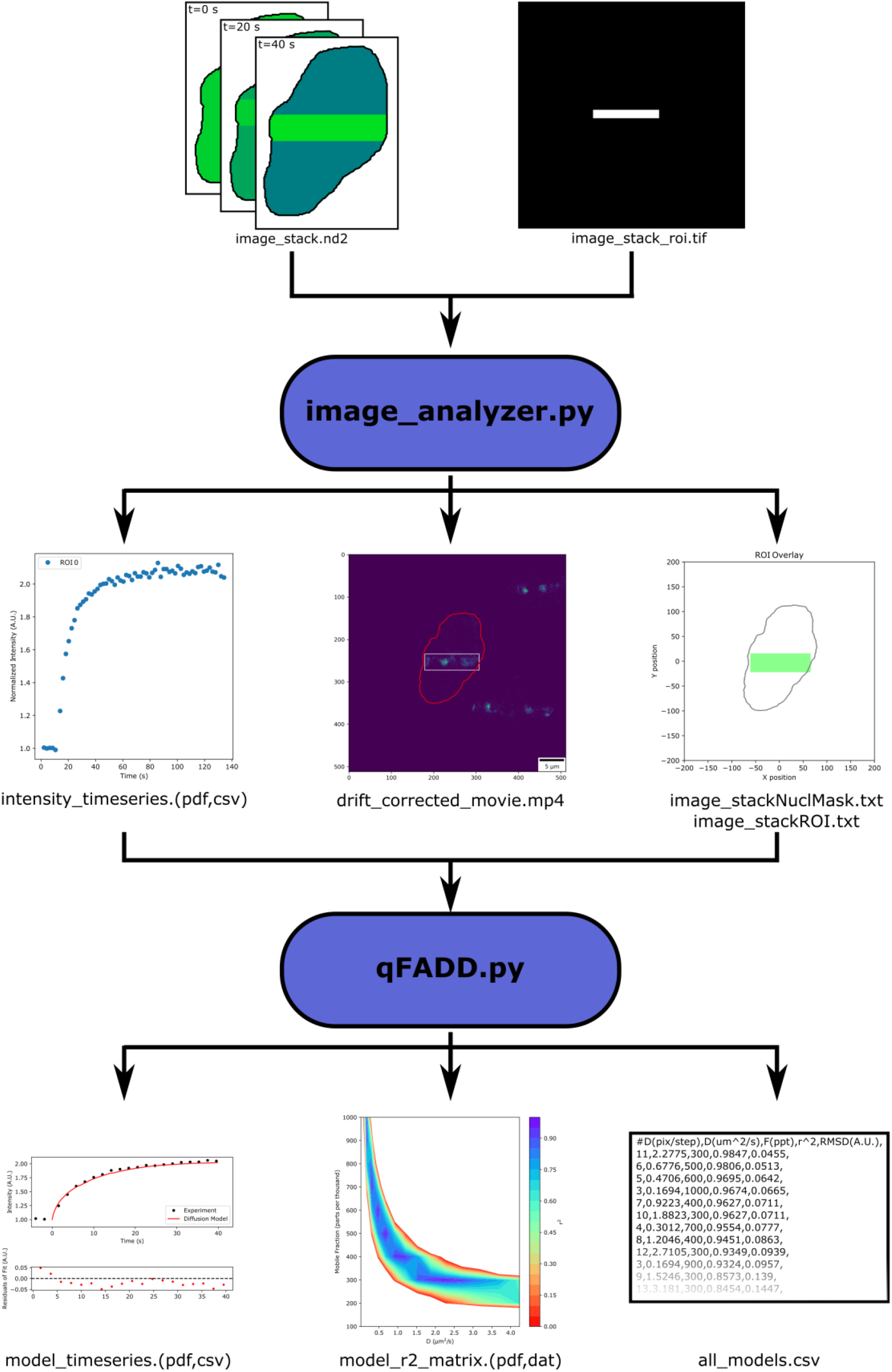
Workflow of the qFADD.py pipeline. Image stacks of fluorescently-tagged protein accumulation and the shape of the region of interest (ROI) are collected by the user on a confocal microscope – here, Nikon/.nd2 files, but all bioformats-readable files are supported. These files are imported into the image_analyzer.py program, which applies motion corrections to the raw image stacks, generates the quantitated accumulation timeseries data, and creates traces of the nuclear envelope and ROI for input to the qFADD.py modeling program. qFADD.py then conducts the grid-search fitting on a range of D_eff_ and ‘mobile fraction’ values defined by the user to identify the best fit model. A plot of the model vs experiment fit is generated, and all sampled models throughout the grid are sorted by fit quality in a human-readable text file (all_models.csv). A visual comparison between models is also provided in a heatmap-style plot of model qualities for rapid assessment of overall performance of the defined grid.

## 2. Methods

Both qFADD.py and associated pre-processing program, image_analyzer.py, are written in Python (v3.6.5), and the code is freely available on GitHub (https://github.com/Luger-Lab/Q-FADD). The publicly accessible Q-FADD server (https://qfadd.colorado.edu) utilizes Anaconda to maintain package distributions, and the stand-alone GUIs (“qFADD_gui.py” and “image_analyzer_gui.py”) are constructed with PySide2 and Qt (v5)(16). Parallel operation is achieved through the mpi4py module, which is constructed on top of the openMPI library(17, 18), and tasks can either be run locally or submitted to a SLURM queuing system(19).

### 2.1. Extraction of Accumulation Timeseries Data

In the laser microirradiation method, fluorescently-tagged proteins are expressed in a cell line of choice. Then, DNA damage is induced in a region of interest (ROI) within the nucleus using a highly focused laser beam, and accumulation of the fluorescently labeled protein(s) in the damaged ROI is tracked. The raw data from these measurements are typically a time-lapsed image stack of the accumulation process and a TIFF image file outlining the ROI, and these are then combined to define the fluorescence intensity timeseries within the ROI.

Previously, the Q-FADD workflow accomplished this through a MatLab notebook that utilized the BioFormats library(20). In the python workflow presented here, this is handled with the “image_analyzer.py” program, which uses the python implementation of BioFormats along with microscope-specific libraries(21). This program takes the ROI TIFF file and the time-lapsed image stack as inputs, and it saves plain text files containing the accumulation data, a mask of the nuclear envelope, and a trace of the ROI to be utilized by the main qFADD.py program.

During this conversion there are several corrections that must be applied in order to maximize the integrity of the accumulation data. First, continuous exposure to the fluorescence excitation laser can cause photobleaching of the fluorophores over the course of the experiment, and the effects of photobleaching must be deconvolved from accumulation and dissipation processes. Therefore, the program applies a scaling factor to the intensity measured at each frame under the assumption that the total number of fluorescent molecules in the nucleus should remain constant:

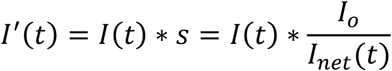

where *I’(t)* is the scaled and outputted ROI intensity value at time *t, I(t)* is the ROI intensity value from the raw image at time *t, s* is the scaling factor, *I*_*o*_ is the frame-averaged total pixel intensities contained within the nucleus in a reference set of frames, and *I*_*net*_*(t)* is the sum of pixel intensities in the nucleus at time *t*. The number of frames to include in the reference set for *I*_*o*_ is declared by the user at runtime, and we elected to reference the first six frames in the example below, as they contained only images of the nucleus prior to the DNA damage event.

A second potential artifact within accumulation data may result from nuclei with lateral drift during image acquisition. In these samples, the nuclear region of DNA damage may migrate beyond the damage ROI expected by the camera, yielding artificially low accumulation counts in later stages of the time course. To correct this error, image_analyzer.py uses the scikit-image library(22), where image translations are identified by determining the cross-correlation between the initial frame and all subsequent frames in Fourier space(23). Notably, this provides the ability to correct for linear motion in the field of view but not for rotations of the nucleus, both in- and out-of-plane with the camera, or for deformation of the nuclear profile. As such, users should still be selective when identifying nuclei for probing and further analysis.

### 2.2. The Q-FADD Algorithm

#### 2.2.1. Nuclear Diffusion Model

In the Q-FADD approach, fluorescence intensity accumulation is modeled using free diffusion in the two dimensional space of the image slice(11). The nuclear mask coordinates are used to determine the boundaries of motion, and movement within the nucleus is conducted on a grid, analogous to the pixel-level information of the acquisition camera. A large number of simulated molecules (here, 10,000) are initially placed randomly along the nuclear grid, and a subset of these molecules – the “mobile fraction” – is selected to participate in the simulated diffusion. Physically, the mobile fraction parameter arises from the fact that some molecules within the nucleus will not move to sites of DNA damage for various reasons.

Molecules within the mobile fraction then undergo diffusion through a Monte Carlo scheme. For every iteration, each mobile particle is able to move either “left” or “right” with equal probability, and the magnitude of the motion is defined by a constant step size per iteration (Δx, in pixels or “grid-points”). The same is true for the motion in the “up” and “down” directions. This results in motion described by:

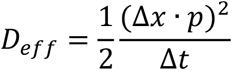

where *D*_*eff*_ is the effective diffusion coefficient, Δx is the step size in pixels per iteration, *p* is the pixel resolution, and Δt is the associated timestep with each iteration. From our experience, Δt should typically be on the order of 0.2 sec per step, or about one order of magnitude higher resolution than the framerate of the camera. However, longer timesteps may be required to model slower diffusing particles, and the loss of sampling between experimental timepoints should be offset with an increased number of replicates per (*D*_*eff*_, mobile fraction) pair. *D*_*eff*_ is used in place of the exact diffusion coefficient as the modeled - and experimentally observed - motion is a convolution of rapid binding and unbinding processes in addition to molecular diffusion.

During the Monte Carlo routine, boundary conditions are applied such that movement beyond the nuclear envelope is rejected. Furthermore, once a molecule has entered the simulated region of DNA damage (defined as the intersection between the ROI of the experiment and the nuclear mask), then it is considered to be trapped at the site of DNA damage. As such, the program is designed to model the accumulation kinetics of damage-response proteins, but it does not account for dissipation of proteins upon DNA repair or completion of signaling task. Users should therefore be careful interpreting results of proteins that dissipate very quickly according to the intensity timeseries.

The simulated trajectory is then converted to an intensity timeseries by counting the number of molecules contained within the ROI at a given iteration of the simulation. To allow for one-to-one comparison between the simulation and experiment, both timeseries are normalized. In this way, I(t) defines the fold-increase of fluorescence (or number of simulated particles) from the initial frame, rather than the exact number of proteins/particles themselves. For the experimental data, normalization is achieved by dividing the ROI intensity of each frame by the average ROI intensity of the pre-damaged frames. In the simulated trajectories, this corresponds to dividing the number of particles in the ROI by the number of particles randomly initialized in the region prior to motion. The simulated intensity timeseries is then interpolated to the experimental timepoints, and fits to data are reported using both the r^2^ metric and the root mean-squared deviation (RMSD) between the theoretical model and the observed fold-increase:

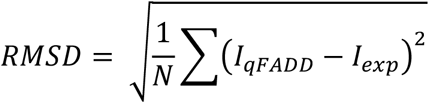

where the sum is carried out over the *N* timepoints in the curve, and *I*_*qFADD*_ is the normalized fluorescence intensity at the interpolated timepoint associated with an experimental normalized intensity, *I*_*exp*_. Both r^2^ and RMSD metrics agree on which parameter combination yields the best model, but the benefits of using RMSD over the canonical r^2^ metric are that RMSD is in the same units of normalized intensity and RMSD can better differentiate models of high fit qualities (Figure 2). The RMSD value thereby tells the user an average discrepancy between the experiment and the model. In this way, RMSD values closer to zero represent high quality fits and poor fits diverge to large values, in contrast to the r^2^ ≈ 1.0 desired in the other metric. However, many users may be more familiar with r^2^ values, so we present the two metrics together both as outputs from the qFADD.py program and here within the manuscript.

**Figure 2.**
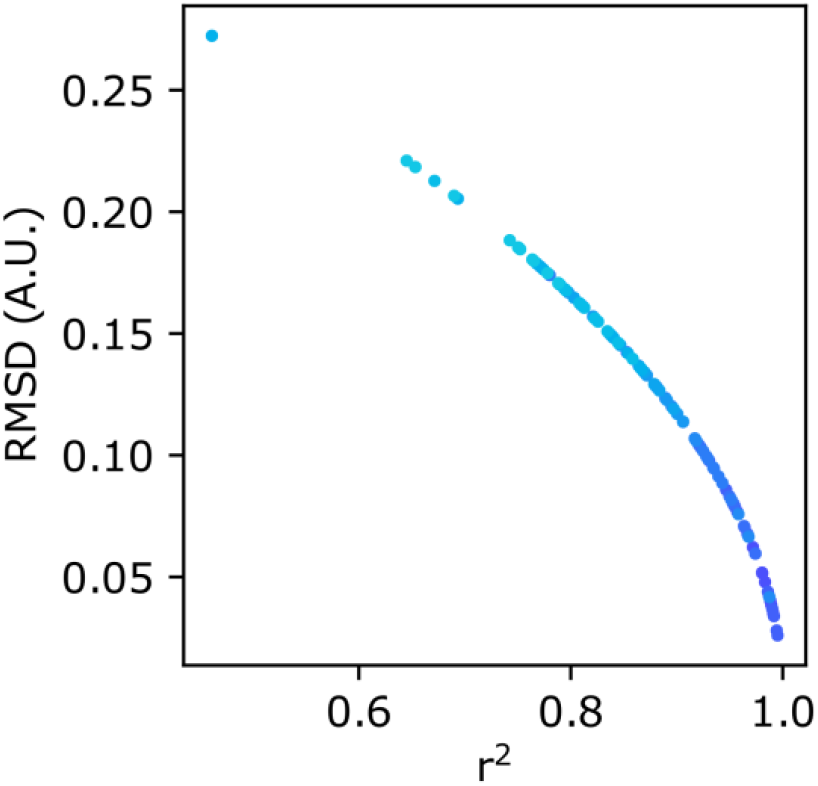
Scatterplot of r^2^ vs RMSD values for individual diffusion models fit to the same nucleus. Each point is color-coded to match replicate simulations using identical (*D*_*eff*_, mobile fraction) combinations (cumulative set of 11 replicas from 10 combinations shown here). While not linearly correlated, r^2^ and RMSD are in perfect agreement for their ranking of each qFADD.py run. Nevertheless, the changing slope to a more vertical line near r^2^ = 1.0 shows that the RMSD metric is better able to differentiate between models of high quality.

#### 2.2.2. Automated Model Fitting

The initial version of the Q-FADD algorithm provided a necessary leap for accurately quantitating repair protein accumulation to sites of DNA damage, namely addressing the relationship between nuclear shape and diffusion behavior. However, it still required users to manually seek out individual combinations of *D*_*eff*_ and mobile fraction through trial and error, which can be difficult and requires large investments of user time and effort. In the new qFADD.py workflow, the task of identifying the model of best fit has been largely removed from the user by utilizing a grid search algorithm. In this approach, users define their desired range and resolution of *D*_*eff*_ and mobile fraction values, and the program evaluates the quality-of-fit for each combination of values. Computational efficiency of this search is enhanced by running tasks in parallel, where each processor is assigned a list of (*D*_*eff*_, mobile fraction) parameter pairs to evaluate. Because Q-FADD is a sampling-based algorithm, multiple replicates are simulated per grid point to ensure that the fit qualities are statistically relevant and not the result of spurious random sampling. As a readout, users can decide between representing the model with either the replica trajectory with the median fit or as an averaged trajectory across all the replicas before fitting to the experiment. With an appropriate number of replicates (*n* > 10 replicates), the model qualities of median and averaged trajectories are typically found to be in good agreement with one another.

While grid search algorithms will identify the single most likely combination of parameters to fit the experimental data, they also produce a variety of alternative models that adequately explain the results. As such, the qFADD.py program both prints the best-fitting model and stores the entire library of sampled parameters and their modeled goodness-of-fit values in an easy-to-read text file. In this way, users can explore the full catalogue of potential models and supplement their intuition gained from orthogonal measurements, rather than fully replacing those results.

### 2.3. image_analyzer.py and qFADD.py Parameters for Example Usage

To demonstrate how to use and interpret the qFADD.py workflow, we provide an example using data gathered upon laser microirradiating mouse embryonic fibroblast (MEF) cells that overexpress the DNA damage signaling protein PARP1, covalently tagged with green fluorescent protein (GFP-PARP1). A full methods description for this experimental set up has been published(11). Cell images were captured on a Nikon A1R laser scanning confocal microscope (Nikon, Tokyo, Japan). Accumulation of GFP-PARP1 was monitored using a 488 nm wavelength argon-ion laser, and DNA damage was introduced with a 405 nm diode laser focused at ∼1.7 mW on a rectangular ROI for 1 sec. Six frames were captured prior to DNA damage for the purpose of normalizing the accumulation timeseries, and the detector resolution was 8.677 nm per pixel. In total, we collected and analyzed data on 11 independent nuclei. One image stack is provided here as an example (example_files.zip), and a modest lateral drift is observed over the course of this collection (Supplemental Movies 1 and 2). This causes the physically damaged region to move slightly beyond the ROI of the camera in the +x direction, meaning that this nucleus may have been manually excluded from the previous Q-FADD workflow, depending on the decision of the user. More drastic motion was observed in a different image stack (Supplemental Movies 3 and 4), and we present this individual nucleus to understand the importance of motion correction in the qFADD.py pipeline (Section 3.1). Presented results were generated using a local installation of qFADD.py, but equivalent results can be gathered by users accessing the webserver (https://qfadd.colorado.edu).

First, the image stack and ROI TIF files were run through image_analyzer.py, where both protein accumulation and the nuclear mask were tracked by the fluorescently-labeled protein channel (“EGFP”). Nuclear masking via the EGFP channel was selected because the protein was overexpressed and localized to the nucleus to provide a quasi-uniform outline of the nuclear boundary, in contrast to the regions of variable brightness observed in the “DAPI” channel. Furthermore, the X- and Y-limits of the ROI were respectively adjusted by −10 and 10 pixels to account for edge effects in the irradiated region, and motion correction was applied. The first six (pre-irradiated) frames were used to normalize the timeseries and correct photobleaching. Then, the nuclear mask and ROI trace files, along with the accumulation timeseries, were fed into a qFADD.py grid fitting which spanned *D*_*eff*_ values of 1 to 15 pixels per iteration and mobile fraction values of 100 to 1000 ppt with a grid point every 100 ppt. Reported fits are the median model from each set of replicates, and 11 replicate trajectories were run per grid point. Each simulation was conducted at a timestep of 0.2 sec per step, yielding a D_*eff*_ range of 0.02 to 4.2 μm^2^/s.

## 3. Results

### 3.1. Effects of Motion Correction

To probe the effects of neglecting motion correction on the measured accumulation timeseries and modeled diffusion behavior, we ran the image stack containing the nucleus with the largest lateral motion through image_analyzer.py with and without motion correction applied, while all other settings were held constant. Indeed, we find that there is a quantitative difference between the raw vs motion corrected timeseries data (Figure 3). The maximum accumulation amount between the two procedures differed by 3.3% (∼2.12 vs ∼2.05 fold accumulation for the motion-corrected and raw data sets, respectively). Moreover, using the timeseries extracted from the raw image stack may lead one to believe that the protein was dissipating from the ROI, starting between 80 to 90 seconds after the damage event (90 to 100 seconds of the collection time). When motion correction is applied, we observe that there is actually no dissipation of GFP-PARP1 from the damage site in this timescale, which was visually confirmed by inspecting the images (Supplemental Movie 3 and 4). While the t_1/2_ values extracted from each timeseries are largely unaffected by the absence of motion correction, their best-fit diffusion coefficients differ by nearly 15% (2.3 μm^2^/s vs 2.7 μm^2^/s for the raw and corrected data, respectively). Together, these data show the importance of properly accounting for nuclear motion during the course of accumulation.

**Figure 3.**
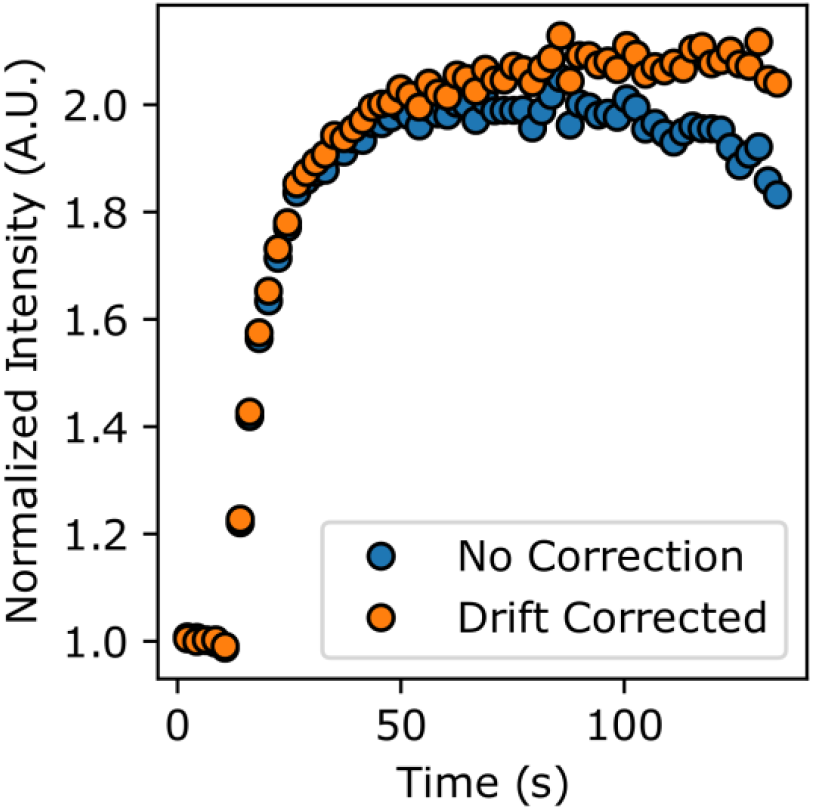
Comparison of accumulation timeseries for raw (blue) and drift corrected (orange) processing. Notably, the curve extracted from the raw movies indicates a dissipation of tagged protein, but properly applying drift correction demonstrates that this is an artifact of the damage site exiting the ROI. Moreover, non-corrected movies show less overall accumulation.

### 3.2. Interpreting Results from a Single Recruitment Image Stack

As an example of the new grid search method, we inspect the qFADD.py modeling results for the nucleus containing a modest amount of lateral motion. This nucleus was selected due to its high fit quality (Figure 4), the presence of several alternative models with appreciably high qualities of fit – which shows the selective power of the grid-search algorithm - and replicates of the same (*D*_*eff*_, mobile fraction) combinations displayed wide ranges of model qualities, as discussed below.

**Figure 4.**
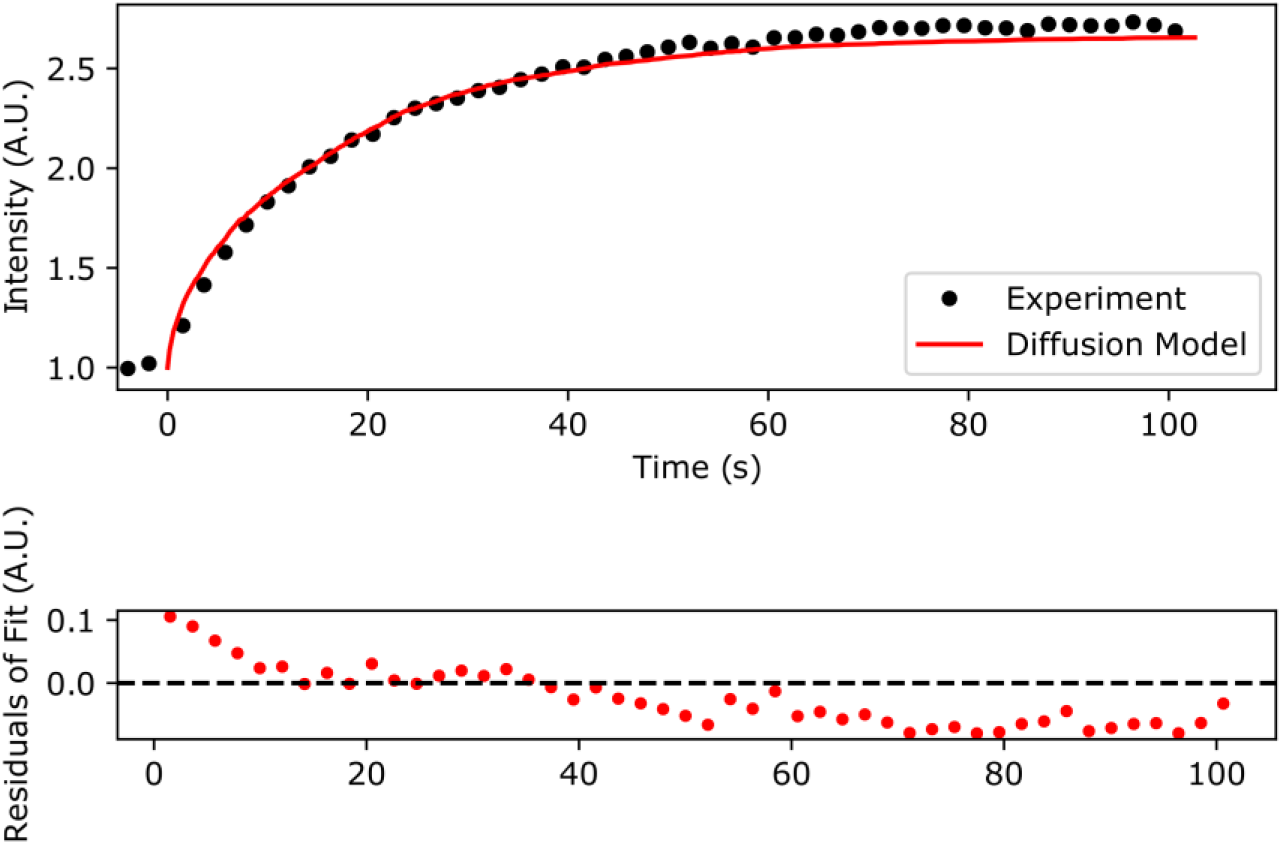
Comparison of the qFADD.py grid search-identified “best fit” model to the experimental accumulation timeseries. (top) Simulated accumulation via free diffusion (red line) vs the experimental profile (black dots). (bottom) Residuals of the fit between the simulation and the experiment. Plotted data are the highest fit-quality median replicate from selected from all sampled median quality replicates in the grid search space (11 replicates per combination of D_eff_ and mobile fraction).

By inspecting the grid search models of the motion-corrected accumulation timeseries (Figure 5A), we find a range of potential models with a mobile fraction of 300 ppt that possess high fit qualities. In this region are the four models possessing r^2^ values greater than 0.9. Two of them have r^2^ values greater than 0.97 (D_eff_ = 11 pix/step, 2.3 µm^2^/s, r^2^ = 0.9807, RMSD = 0.0515 and Deff = 10 pix/step, 1.9 µm^2^/s, r^2^ = 0.9742, RMSD = 0.0596). These data show that multiple combinations of D_eff_ and mobile fraction can achieve acceptable fits, depending on the chosen definition of “acceptable” r_2_ value. In this way, the grid-search method implemented in qFADD.py robustly identifies the model of optimum fit by scanning a wide range of values and running multiple replicates per parameter combination, whereas a user searching for the best-fit model via trial and error may have been satisfied with identifying one of the suboptimal model fits, if they were discovered before the optimal fit.

**Figure 5.**
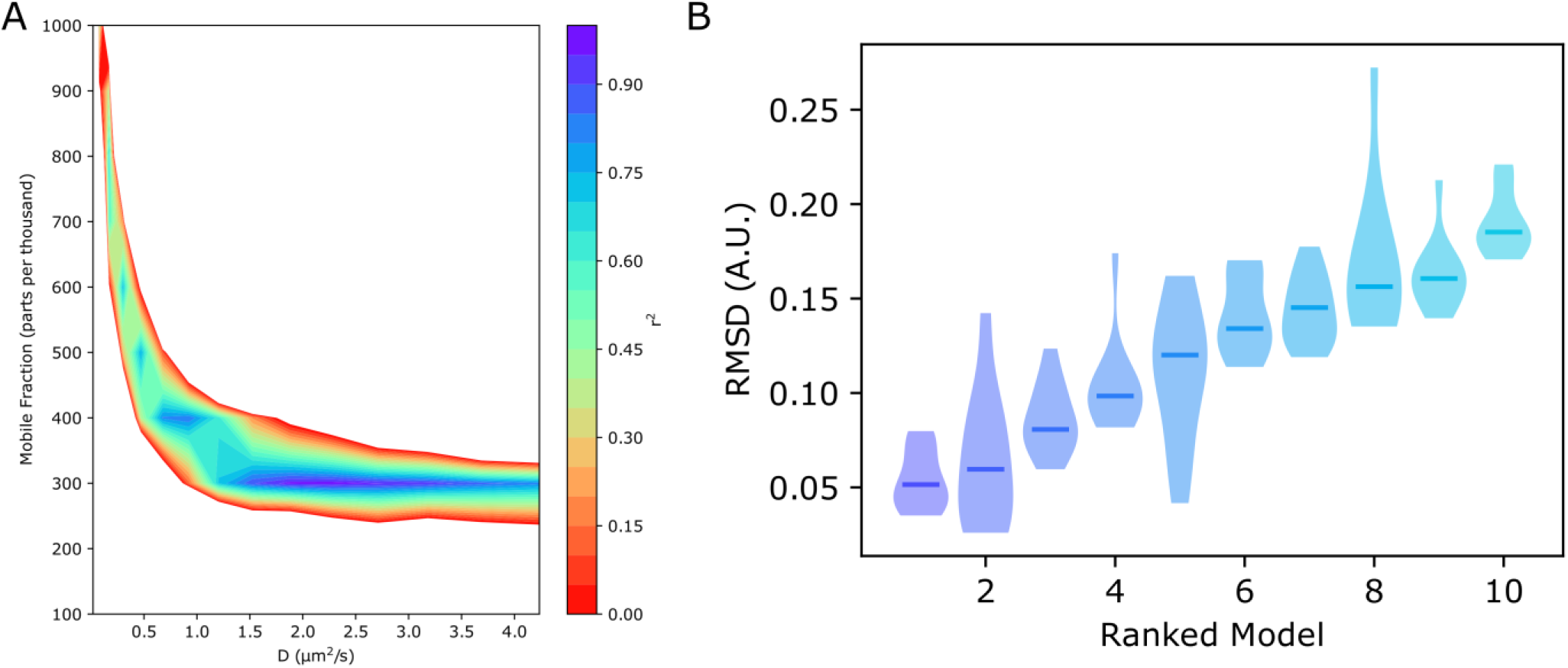
Comparisons of sampled model performances during the qFADD.py grid-search algorithm. (A) Heatmap showing multiple local regions of relatively high (r^2^ > 0.9) fit, showing that several different parameter sets may adequately describe the experimental profile, depending on user-definition of high-fidelity. However, only the population centered around a mobile fraction of 300 ppt contains the models with r^2^ values greater than 0.95. (B) Violin plots showing the RMSD distributions across the 11 independent replicates for the ten best performing (*D*_*eff*_, mobile fraction) combinations on the same experimental data set. Lower values correlate to simulated models that are more similar to the experimental profile. These plots show that some parameter combinations have spuriously high (or even low) fit qualities, whereas their median fit qualities (solid lines) provide a more robust ranking metric

To emphasize this point, we inspect the model qualities of the 11 replicates sampled during the qFADD.py grid-search (Figure 5B). In this search, the first replicate identified with an r^2^ value greater than 0.95 is the (*D*_*eff*_, mobile fraction) combination of 9 pix/step (1.5 μm^2^/s) and 300 ppt (RMSD of 0.0758 from the experimental profile). In the previous Q-FADD pipeline, a user satisfied with a r^2^ > 0.95 fit quality might accept this model and not consider additional simulations. However, our grid-search results show several better performing models, and the (*D*_*eff*_, mobile fraction) combination of (9 pix/step, 300 ppt) is only the 5^th^ best model identified when considering the full array of replicates (median r^2^ of 0.8952, median RMSD of 0.1201). For the best model identified during the grid search (D_eff_ = 2.3 µm^2^/s, mobile fraction = 300 ppt), we find that the r^2^ values range from 0.9572 to 0.9910 (RMSD range of 0.0767 to 0.0351). Furthermore, the parameter combination with the best single replicate (D_eff_ = 1.9 μm^2^/s, mobile fraction = 300 ppt) is the only the second-best model overall, with an r^2^ of 0.9951 and RMSD value of 0.026. In addition to the median overall fit quality, the worst-fit replicate from this parameter combination (r^2^=0.8529, RMSD=0.0788) is notably worse than that of the selected “best-fit” model, and the distribution of fit qualities across this second-best state is much wider than the best identified parameter set. Indeed, if users had used a single replicate of any set of parameters, they may have identified the best-fit model to possess a D_eff_ of 9 pix/step (1.5 μm^2^/s), which would have lead to a discrepancy of 33% with the more statistically robust D_eff_ of 11 pix/step (2.3 μm^2^/s) model. Similarly, using a trial-and-error approach with single replicates may lead a user to consider the D_eff_ = 10 pix/step (1.9 μm^2^/s) to be a rather poor fit to the data (r^2^ < 0.9), despite being the second most competitive model, and thus may have caused them to drastically alter either their mobile fraction or D_eff_ values away from the grid search-identified global maximum fit-quality space. This bias away from the “correct” model can yield a substantial increase in user effort and the active time spent searching for a model of appreciable quality, but the grid-search employed within qFADD.py efficiently protects users from such results, as the automated search reports on a wide number of parameter combinations while simultaneously requiring no interaction from the user after the limiting values of each parameter combination are defined.

### 3.3. Combining Results from Multiple Nuclei

Above, we have demonstrated the functionality of the qFADD.py grid-search approach conducted on a single nucleus. However, one of the strengths of the Q-FADD approach is the ability to identify a distribution of D_eff_ and mobile fraction values observed across a variety of nuclei, regardless of their shapes and sizes, rather than extracting a singular kinetic rate from the averaging of data from many differently shaped nuclei. To this end, both the qFADD.py and image_analyzer.py programs can be run in a “batch” mode, where a collection of nuclei can be simultaneously analyzed. In practice, users wishing to utilize the “batch” processing mode should be cautious that every nucleus in the dataset possesses identical parameters, such as number of normalization frames prior to irradiation, the time of the irradiation event, and that images were collected through the point of maximum accumulation and not truncated while recruitment was still ongoing. We also provide the “qfadd_distribution.py” program, which builds violin plots from a cumulative set of Q-FADD results (Figure 6A), and this program is also capable of comparing models across multiple data sets (Figure 6B,C).

**Figure 6.**
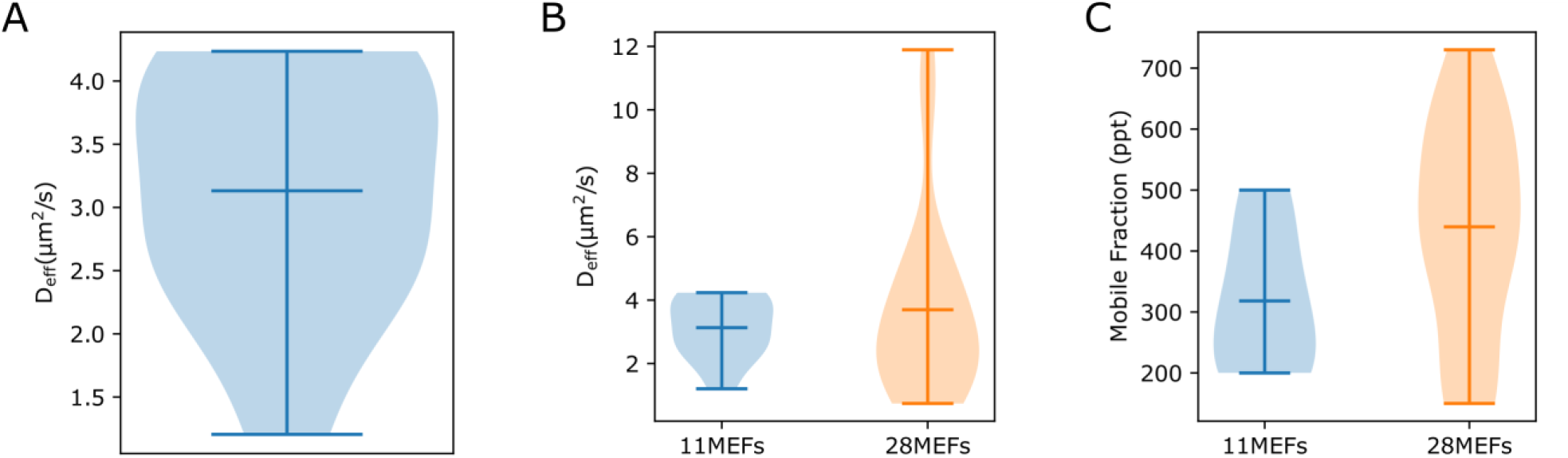
Examples of the comparison violin plots that can be generated with the qfadd_distribution.py program. (A) Plot of D_eff_ values generated when running the program on a single collection of models (here, GFP-PARP1 overexpressed in 11 independent MEF cells). The shaded regions define the relative frequency distribution of reported D_eff_ values (wider regions correspond to more frequently observed D_eff_ values), and the horizontal lines define the maximum, mean, and minimum values observed. (B) Plot of D_eff_ generated when comparing between multiple datasets. “11MEFs” represents the 11 cells sampled in the work of this manuscript, and the “28MEFs” dataset is from our previous publication (11). While the models from the larger data set (28MEFs) contain a wider range of D_eff_ values, the two distributions are in agreement (p-value of 0.36, also reported by the qfadd_distribution.py program). (C) Comparison plot of mobile fraction values, which provides a notably wider distribution than the D_eff_ plots of either collection set.

To showcase this functionality, we collected microirradiation measurements on 11 separate nuclei, including the previously described example, and we inspected the range of diffusion coefficients reported by the 11 best-fit models identified through our qFADD.py grid search (Figure 6A). Modeled D_eff_ values range from less than 1.5 µm^2^/s to greater than 4 µm^2^/s, showcasing the wide variability observed in microirradiation data as a result of nuclear shape and size, even on samples from the same collection session. This distribution was then compared to the model values previously reported by our manual-search Q-FADD pipeline conducted on 28 MEF cells overexpressing GFP-labeled PARP1 (Figure 6B). This comparison shows that there is no significant difference between the modeled diffusion rates in the two data sets (D_eff_ = 3.1 ± 0.3 vs 3.2 ± 0.3 µm^2^/s, p-value of 0.36), but a modestly significant difference was noted in the mobile fraction comparison (318.2 ± 33.6 ppt vs 439.7 ± 33.3 ppt, p-value of 0.02, Figure 6C). The mobile fraction distribution is notably more wide-spread than the D_eff_ distribution of either data set, suggesting that this discrepancy may be a result of collecting fewer nuclei than was conducted previously, but this disparity could also be explained by a number of other factors, such as location of sampled cells within the cell cycle, resolution differences between the “coarse-grained” grid-searched mobile fraction (here, 100 ppt resolution) vs the “high-resolution” trial-and-error search conducted previously (1 ppt resolution), lack of motion correction in the original dataset, or potential user selection of sub-optimal models from benevolently sampling high-quality fits from individual parameter combinations.

## 4. Discussion

Here, we have described the qFADD.py analysis pipeline, an open-source Python implementation of the Q-FADD method(11) with updated image processing, diffusion-fitting procedures, and additional analysis tools. The software is available as both a standalone program for local installation (https://github.com/Luger-Lab/Q-FADD) and through a publicly accessible webserver (https://qfadd.colorado.edu). By providing these two integrations, researchers with modest computing resources can analyze their microirradiation data via the webserver, while labs with cluster access can make efficient use of the parallelized grid search algorithm on their local resources.

The qFADD.py pipeline has introduced several updates to the original Mathematica and MatLab versions. One major improvement is the addition of nuclear tracking to the image processing workflow, which corrects for lateral drift and allows researchers to avoid discarding valuable image stacks, as well as safeguarding them from interpreting nuclear drift as the dissipation of fluorescently-labeled proteins from the induced damage site. This allows researchers to maximize the number of usable data sets in their analysis and improves statistical rigor when comparing accumulation kinetics between cell conditions or proteins of interest. We have shown here that modeling accumulation kinetics from non-corrected image stacks can lead to quantifiable differences in predicted values. Furthermore, the timeseries data extracted by the image_analyzer.py program is not limited to use only in the qFADD.py program, but it can also be used to extract dissipation timeseries for modeling parameters such as off-rate kinetics. As such, motion correction applies another layer of protection from incorrectly fitting off-rates to false dissipation events, but the corrections applied in image_analyzer.py still have some limitations. Notably, the algorithm only corrects for lateral drift and cannot account for cells that undergo rotations either in-or out-of-plane with the camera. Additionally, nuclei that significantly deform due to continuous exposures with intense light cannot currently be corrected. In addition to the development of 3-dimensional nuclear mapping, these limitations are the predominant geometric constraints on the Q-FADD pipeline.

While motion correction during image processing extends the number of nuclei that can be included in final datasets, careful selection of nuclei should still be performed while collecting microscope data. Nuclei with blebs may pose issues for accurately tracing the nuclear envelope, and these nuclei may subsequently yield D_eff_ and mobile fraction combinations that can satisfy the experimental curve but with an improper nuclear geometry, thereby representing a false-positive result. Additionally, users should carefully control microscope temperature and humidity in order to maintain healthy cells over the course of collections.

The second key improvement to the original Q-FADD algorithm is the utilization of the automated grid search to identify the best-fit model from a collection of potential models. With the use of multiple replicates per parameter set, this method determines the best-fit model in a statistically robust way, whereas a user’s trial-and-error search may be prone to identifying spuriously high-fit models. On the other hand, the grid search method requires a balance of resolution and computational efficiency, as searching out a wide range of mobile fractions on the single part per thousand level is exceptionally demanding. Instead, users should first identify general regions of high model fidelity utilizing a coarse grid step (such as 100 ppt), and then focus in on the boundaries of said regions at higher resolution (such as 10 or 25 ppt) in order to extract more precise values of this metric. However, the resolution limit of the mobile fraction parameter is largely dependent on the recruitment speed, protein expression levels, and fluorescent labeling efficiency, so users should be careful to avoid overfitting the mobile fraction value with single part per thousand resolution in systems where 10 to 50 ppt will suffice.

With its automated workflow behavior, the qFADD.py pipeline provides researchers with a straightforward way to model the kinetics of proteins that operate in a variety of DNA repair pathways. In combination with multi-photon approaches, this method allows the opportunity to simultaneously probe the dynamics of several proteins within the same nucleus, rather than relying on separate image stacks from different nuclei.

## Supporting information

example_files.zip

## 5. Author Contributions

S.B., J.M., and J.R. designed the updated qFADD.py and image_analyzer.py programs and outputs. S.B. coded both programs, and P.B. provided code blocks for the qFADD.py program. S.B. and J.M. designed the stand-alone GUIs, and S.B. constructed them. P.B. and J.M. conducted the cell experiments and image capturing. S.B. generated and interpreted the computational results. K.L. secured funding for the project. S.B. wrote the original draft of the manuscript, and all authors contributed to its editing.

## 6. Acknowledgements

The authors thank the CU Boulder BioFrontiers Institute IT department, in particular Ethan Kern, Matthew Hynes-Grace, and Jon Demasi, for their assistance with implementing the publicly-accessible webserver. We also thank Drs. Jian Wey Tay and Erik Grumstrup for providing the original MatLab and Mathematica codes for porting to Python, as well as Asmita Jha for providing feedback on the user experience of both the webserver and standalone versions of the program.

J.M., J.R., and K.L. are funded by the National Institutes of Health – National Cancer Institute (R01 CA21825), J.M is also funded by the American Heart Association (20POST35211059), and J.M. and K.L. were further supported during this work by the University of Colorado Cancer Center (Pilot Funding Grant ST63501792). Members of the Luger Lab are also supported by the Howard Hughes Medical Institute.

## Notes

### Competing Interest Statement

The authors have declared no competing interest.

