## Supplementary material for "Automated Modeling of Protein Accumulation at DNA Damage Sites using qFADD.py": example_files.zip: qFADD_gui_documentation.pdf

### How to Use qFADD.py Pipeline GUIs

The qFADD.py pipeline is broken down in to two steps. First, you use a collection of image stacks and a .tif file outlining the damage ROI (from the microscope) to trace the nuclear envelope (the nuclear mask) and extract the fluorescence intensity timeseries to the region of damage. Then, you use the map of the nuclear envelope and the ROI definition, in conjunction with the extracted fluorescence intensity timeseries, to fit an array of Monte Carlo-based diffusion models to the experimental timeseries and extract the model of best-fit from an automated grid search.

#### 1. The Image Analyzer (pre-processing)

1. Start the Image Analyzer interface by calling "image_analyzer_gui.py" from the command line.

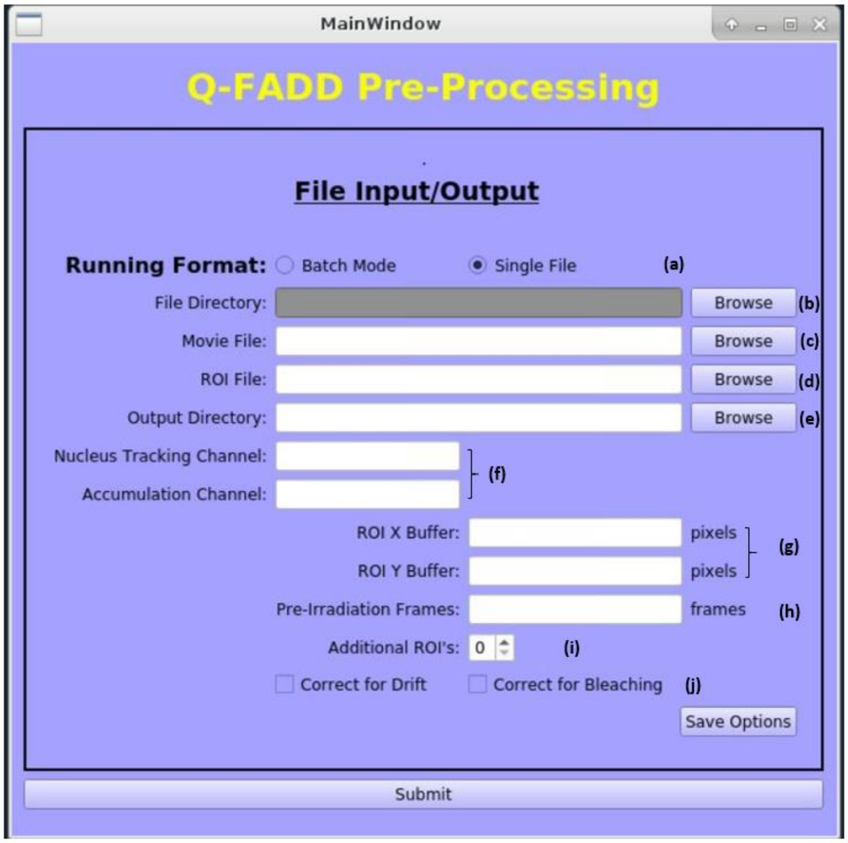


***Figure 1. The GUI for the Image Analyzer pre-processing program.*** *The program runs on both single files and batches of files contained within the same folder. A full description of each option can be found below.*

2. The Image Analyzer has several adjustable parameters, as follows:
   (a) Running Format: The pre-processing routines can either be run on the image stack and ROI from a single acquisition ("Single File" mode) or on a collection of acquisitions stored in the same directory ("Batch Mode").
      - Because "Batch Mode" will process all files identically, it is important that certain acquisition parameters (such as the number of frames collected prior to damage, as well as the channel IDs) are identical across the collections.

(b) File Directory: When "Batch Mode" is selected, this option will define the location of the directory that houses all the files that you wish to process. In "Single File" mode, this field is not used by the program, and is "greyed-out".

(c) Movie File: This input tells the program how to identify the acquisition stack from which the accumulation profile will be extracted.
- When "Single File" mode is selected, you can either type the path to where the image stack is located, or you can click the "Browse" button to find it through the file manager GUI.
- In "Batch Mode", this setting is replaced with "Movie File(s) Extension", and you need only to type the extension by which you can distinguish the image stacks in the directory from any other files in the directory (for example, ".nd2").

(d) ROI File: This input tells the program how to identify the ROI-definition file that should be linked to the image stack(s).
- When "Single File" mode is selected, you can either type the path to where the ROI .tif file is located, or you can click the "Browse" button to find it through the file manager GUI.
- In "Batch Mode", this setting is replaced with "ROI File(s) Extension" and is used to distinguish ROI files from other types of files in the directory (for example, "_ROI.tif").
  - To link a particular file with the "_ROI.tif" extension with the associated ".nd2" file, the filenames must match up until the different extensions. For example, "file1.nd2" and "file1_ROI.tif" will be linked, but "file1.nd2" and "file_1_ROI.tif" will not.

(e) Output Directory: This defines the location where the output files will be sent after analysis is complete. This can be a new or pre-existing directory.
- Since you may run several different datasets that need to be compared, such as acquisitions in the presence of inhibitors at various concentrations, please be sure to use directory names that will help you identify each data set.

(f) Nucleus Tracking and Accumulation Channels: These help the program identify which imaging channel to use for identifying the nuclear envelope ("Nucleus Tracking") and for extracting the accumulation profile ("Accumulation").
- The "Nucleus Tracking" channel should be the channel that best traces the nucleus of your system. For some datasets, this may be the DNA-stain channel (i.e., "DAPI") but for others, this may be the labeled-protein channel (i.e., "EGFP").
  - Depending on the imagestack filetype (i.e., ".nd2" vs ".tif" files), channel names may be strings, as listed above, or they may be integer identifiers, such as "0", "1", or "2".

(g) ROI Buffer: These fields tell the program how much to pad/trim the ROI region by, to avoid loss of accumulation pixels due to edge effects of the damage laser. In our hands, "-10" in the X-dimension and "10" in the Y-dimension work well, but you may wish to explore these values more on a imagestack-by-imagestack basis.

(h) Pre-Irradiation Frames: This is the number of pre-damage event frames recorded for your image stack(s) and is used to normalize the intensity timeseries. For "Batch Mode", it is imperative that this value is the same across all Movie Files.

(i) Additional ROIs: If this value is non-zero, the Image Analyzer will generate intensity kinetics data for this many ROIs above and below the damage site, to track the depletion of labeled-protein from those regions.

(j) Correct for Drift and Bleaching: These checkboxes tell the program to run corrections to the image stacks before exporting the accumulation timeseries, and they should typically both be checked for every run.

- Drift correction is necessary to distinguish between bonafide dissipation events and artificial depletion of intensity due to a nuclear damage site drifting beyond the expected ROI.
- Bleach correction will remove the effects of photobleaching caused by repeated exposure to the excitation laser over the timecourse.

- If the same settings will be used repeatedly, you may save them by clicking "Save Options" to make them your default parameters, but this step is not required for the analysis to run on your given inputs.
- Click "Submit" to start the analysis.
    - Due to some limitations in how the GUI initializes the javabridge tasks that integrates with the movies, the Image Analyzer GUI must remain open over the course of the analysis.

**Once the analysis is complete, the following data files will be generated:**

i. Two Intensity kinetics files (.csv) containing a list of timestamps (s) and intensity values (one file containing the raw values and the other containing normalized values) for the requested ROIs.

ii. Two video files (.mp4) showing the tracking of the nucleus (raw and drift corrected).
   a. In these video files, the ROI is outlined with a white border and the nuclear envelope is traced by a red line.
   b. If your tracing is quite poor, you should first try using a different channel that may better identify the edges of the nucleus without additional features inside.
      i. If your tracing problems still persist, please contact us using the "Issues" tab of the GitHub repository: https://github.com/Luger-Lab/Q-FADD/issues

iii. A plot of normalized intensity timeseries (.pdf)
   a. If this plot does not look like the expected accumulation behavior, then please closely inspect the raw and drift corrected .mp4 files to ensure that your nuclear tracking is sufficient and that the damage site indeed spends the timecourse within the ROI.
   b. If this timecourse has a very weak signal-to-noise ratio, then it may be difficult to robustly fit a model to the data using Q-FADD. Therefore, you may want to consider altering your experimental setup to increase the fluorescence signal, or simply omit this nucleus from further analysis (if there is an appreciable number of other nuclei with better signal-to-noise).

iv. A file (.txt) consisting of the coordinates for the nuclear mask.

v. A file (.txt) consisting of the coordinates for the ROI.

### 2. The Q-FADD Grid-Search (Diffusion Modeling)

1. Start the Q-FADD interface by calling “qFADD_gui.py” from the command line.

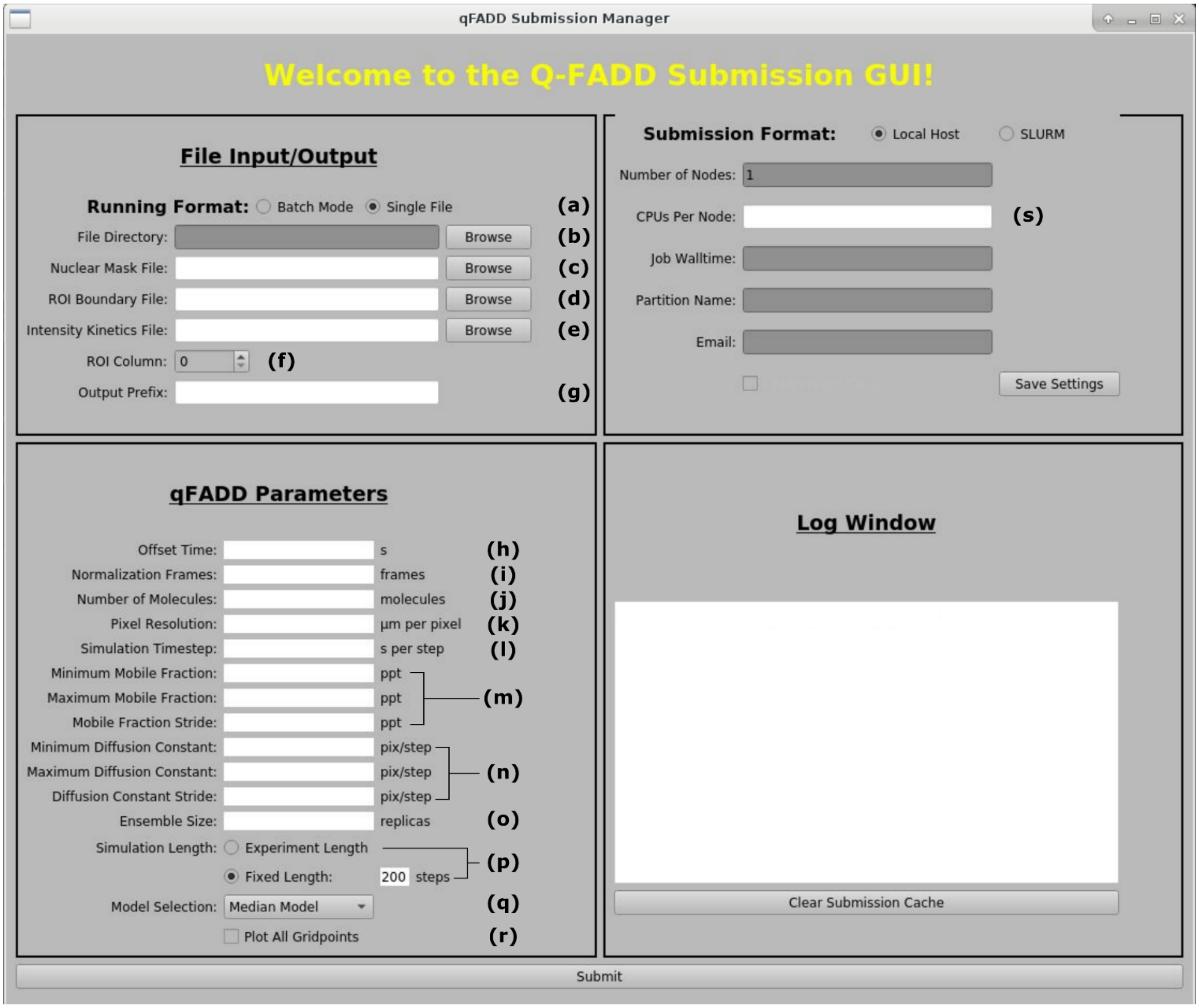


***Figure 2. The Q-FADD graphical interface.*** *The interface is separated in to four different sections: File Input/Output, qFADD Parameters, Submission Format, and the Log Window. A full description of the parameters can be found below.*

2. The File Input and Output parameters are as follows:
   (a) <u>Running Format:</u> Similar to the Image Analyzer pre-processing, qFADD.py can generate Q-FADD models either in “Single File” or “Batch Mode” schemes.
   - Unlike the image analysis routine, which should have identical parameters if the experiments were set up carefully (making it quite easy to run in batch mode), Q-FADD directly accounts for variations between nuclei, and therefore identical parameters may not fit every nuclei, as noted below.

   (b) <u>File Directory:</u> This parameter is only used in “Batch Mode”, and it identifies the folder containing the collection of Nuclear Mask, ROI Boundary, and Intensity Kinetics files.

(c) Nuclear Mask File: This identifies the nuclear mask .txt file (that was generated by the Image Analyzer pre-processing) that provides the coordinates for the nuclear envelope.

- In "Batch Mode", this option is changed to "Mask Extension", and the extension of nuclear mask-type files within the "File Directory" should be provided (typically "NuclMask.txt", unless these files have been renamed by the user).

(d) ROI Boundary File: This identifies the ROI coordinate .txt file (generated by the pre-processing program) that provides the region within the nucleus that tagged molecules will be "trapped" on DNA damage for the nucleus outlined by "Nuclear Mask File".

- In "Batch Mode", this option is changed to "ROI Extension", and the extension of ROI-type files within the "File Directory" should be provided (typically "ROI.txt", unless these files have ben renamed by the user).

(e) Intensity Kinetics File: This identifies the file that contains the fluorescence intensity accumulation timeseries .csv file associated with the input "Nuclear Mask File".

- In "Batch Mode", this option is changed to "Intensity Kinetics Extension", and the extension of the intensity kinetics files within the "File Directory" should be provided (typically ".csv" or "_normalized.csv", unless modified by the user).

(f) ROI Column: This is the column (0-indexed) within the "Intensity Kinetics File" that you would like to fit with Q-FADD diffusion models.

- For example, in the figure below, if we were interested in fitting the EGFP ROI 0 values, then we would enter "2" for the "ROI Column" because the $0^{th}$ column is the time, the $1^{st}$ column is the "DAPI – ROI 0" values, and the $2^{nd}$ column is the "EGFP – ROI 0" values.

| | DAPI | EGFP |
|---|---|---|
| Time (s) | ROI 0 | ROI 0 |
| 0.083343 | 757560 | 382650 |
| 2.189361 | 767720 | 379945 |
| 4.267277 | 770506 | 376465 |

***Figure 3. Fraction of the Intensity Kinetics File that helps us determine "ROI Column" value.***

(g) Output Prefix: This provides a unique prefix to the outputs generated from this run.

- Users should be define a unique prefix so that they can quickly identify different fitting protocols that they may have used on each nucleus.

3. The qFADD paramaters are as follows:

(h) Offset Time: This is the timepoint (in sec) at which the damage is introduced and the repair protein starts to accumulate.

- When running in "Batch Mode", this parameter may vary slightly between nuclei (and is potentially influenced by changing ROI shape and size between nuclei in a single collection), and can be the cause for poor fits across the batch. Therefore, users should visually inspect poor fits from the batch processing and run them again in "Single File" mode, with altered parameters.

(i) Normalization Frames: This is the number of pre-irradiation frames, used to normalize the experimental timeseries. Should be identical to the value used in the pre-processing Image Analyzer.

(j) Number of Molecules: This is the number of simulated particles that will be placed within the nucleus. In our hands, values of ~10,000 work well to establish good signal-to-noise in the sampling algorithm.

(k) Pixel Resolution: This is the resolution of the initial image stacks (in µm per pixel).
  - Because the simulations are run at the pixel-level resolution for 1-to-1 comparison with the image stack, it is imperative that all files in the "Batch Mode" processing scheme have the same pixel resolution.
  - While data from image stacks using different pixel resolutions cannot be run in "Batch Mode", it is perfectly acceptable to group the final modeling results (in terms of $D_{eff}$ and mobile fraction values) into the same final distribution of values.

(l) Simulation Timestep: This value sets the number of seconds per iteration of the Monte Carlo diffusion simulation, and it is used to convert the diffusion constant from pixels/s to $\mu m^2/s$. Lower timestep values correspond to faster effective diffusion constants.
  - To ensure good temporal sampling, this value should typically be an order of magnitude lower than the timestep between experimental frames.
  - In the cases of extremely slowly diffusing molecules, it may be necessary to increase the simulation timestep to ½ the resolution of the experiment.
    - Because larger values of "Simulation Timestep" corresponds with less efficient temporal sampling, users that increase their "Simulation Timestep" should similarly increase their "Ensemble Size" to counteract the lost sampling efficiency in determining statistical robustness.

(m) Mobile Fraction (Minimum, Maximum, and Stride): These values set the limits (and step size from minimum to maximum) for the "mobile fraction" dimension of the grid search.
  - Because the total number of Q-FADD simulations is (Number of Mobile Fraction Values) x (Number of Diffusion Constant Values), excessive precision in this metric can cause a rapid increase in the computational cost of the grid search.
    - To most efficiently sample the ($D_{eff}$, mobile fraction) phase space, users are recommended to do an initial grid search across a wide range with low precision ("Stride" ≥ 100 ppt), and then focusing in with subsequent Q-FADD models that span regions of high fit quality with a narrower minimum to maximum range, but with higher precision (lower "Stride" values).

(n) Diffusion Constant (Minimum, Maximum and Stride): These values set the limits (and step size from minimum to maximum) for the "Diffusion Constant" dimension of the grid search.

(o) Ensemble Size: This is the number of simulated replicates per ($D_{eff}$, mobile fraction) grid point. In our hands, 11 or more replicates produce the greatest statistical robustness when ranking the sampled models.

(p) Simulation Length: There are two ways to determine the simulation length:
  - *Experiment Length:* Set the simulation length to match the time for maximum accumulation in the experimental profile.
  - *Fixed Length:* Alternatively, users can define the number of iterations (steps) in each Monte Carlo diffusion model. The total simulated time is then (Number of Steps) x (Simulation Timestep).

(q) Model Selection: Users can choose between two methods for selecting a representative model per grid point. Because Q-FADD is a sampling-based technique, spuriously high or low performing models will influence model ranking, so these choices are designed to improve statistical robustness in ranking different ($D_{eff}$, mobile fraction) parameter sets.

- *Median Model:* For each ($D_{eff}$, mobile fraction) combination, the reported model for ranking is the model from the ensemble with the median quality-of-fit value with the experiment.
    - If "Median Model" is selected, users should simulate an odd number of replicates.
- *Average of Models:* With this selection, qFADD.py will average the accumulation timeseries of all replicates for a single parameter combination, and then it will rank all parameter combinations according to the quality-of-fit for this mean timeseries.

(r) <u>Plot All Gridpoints:</u> Checking this box will cause the program to generate a PDF figure for the simulated vs experimental timeseries in all parameter combinations, rather than only the best-identified model.

4. Submission Format parameters:

(s) <u>CPUs per Node:</u> This value determines how many processors the Q-FADD simulations will be spread across.

- Users have the option to run the qFADD.py grid search on their local machine, or to submit the task through a SLURM-style queuing system. This will allow access to many CPUs, in parallel, which can help users simulate wider grid ranges with higher precision (lower "stride" values). However, because SLURM systems are not uniformly defined, the values for these parameters should be determined with the assistance of the user's technical support staff.

**Once the analysis is complete, the following files will be generated:**

- Each of these files will be located within a folder that takes the file name of the nucleus, but the files themselves will be prefixed by the value you supply to (g) "Output Prefix"

i. A text document that lists the parameters entered for the simulation ("_inputs.par").

ii. A comma-separated values (.csv) file that contains all sampled models and performances, ranked from highest to lowest performing models ("_all_models.csv").

iii. $r^2$ performance metrics for the sampled models are shown in a matrix-format text file ("_r2_matrix.dat") as well as in a PDF-format heatmap figure ("_r2_matrix.pdf"), for quick assessment of grid-search performance.

iv. The same information, but using the RMSD assessment metric, is also provided ("_rmsd_matrix.dat" and "_rmsd_matrix.pdf"). Qualitatively, these two plots will typically be in high agreement with one another, but the RMSD metric provides better separation between models of high fitness quality.

v. A PDF-format figure that shows the intensity vs time plots of both the experimental and simulated profiles, along with the residuals of best fit ("_Intensity_Timeseries.pdf").

vi. A comma-separated values (.csv) file that contains the simulated and experimental timeseries values ("_roi0_intensity_timeseries.csv").

vii. A PDF-format figure that shows the nuclear envelope and overlapping ROI definition ("_Model_Nucleus_with_ROI.pdf").
