## Supplementary material for "Automated Modeling of Protein Accumulation at DNA Damage Sites using qFADD.py": example_files.zip: QFADD_server_tutorial.pdf

### Using the Q-FADD Server

This document outlines the basic workflow for using the Q-FADD webserver. Default values are provided by the webserver for many parameters, but users should be aware that the best parameter for their own modeling efforts may differ greatly from the default values, depending on the experimental design and diffusive behavior of the protein of interest. For questions or concerns with workflow usage, performance, or results, please use the "Issues" feature of the qFADD.py GitHub repository: [https://github.com/Luger-Lab/Q-FADD/issues](https://github.com/Luger-Lab/Q-FADD/issues).

#### 1. The Image Analyzer

Before diffusion models can be fit to the experiment, fluorescence intensity timeseries data and a map of the nuclear envelope must be constructed. This is achieved through the "Image Analyzer" module of the webserver. A snapshot of the Image Analyzer interface is shown below.

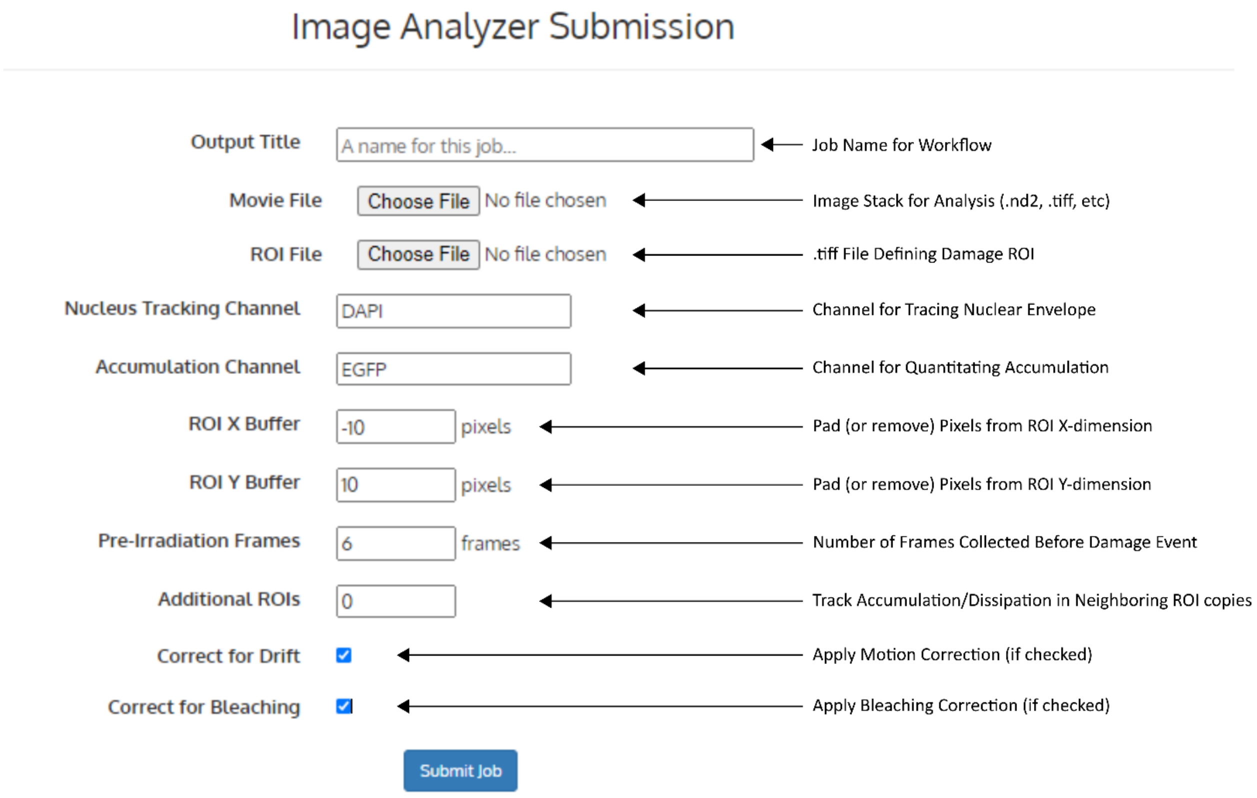


***Figure 1. Snapshot of the Image Analyzer interface.*** *Movie and ROI files are uploaded from the user's local computer, and output files are stored on the Q-FADD server for download.*

Input Parameters:

- Output Title: This is the name for the workflow, as will be shown in the “Pipeline Results” tab of the webserver.
- Movie File: Users upload their image stack from their local computer. Currently, full support is provided for Nikon .nd2 files, and limited support is offered for all other bioformats-supported filetypes.
- ROI File: A .tiff file that outlines the damage region-of-interest (ROI). In the current version, only rectangular ROIs are supported.
- Nucleus Tracking Channel: This is the name of the optical channel from (Movie File) that will be used to determine the nuclear shape and correct for beam-induced motion. Depending on the filetype of (Movie File), this could be a string (i.e., “EGFP”) or an integer channel ID.
    - For best tracing of the nuclear envelope, users should select the channel that best outlines the nuclear envelope, which may in some cases be the protein-label channel and not the DNA-stain channel.
- Accumulation Channel: Similar to the (Nucleus Tracking Channel), this identifier tells the Image Analyzer which optical channel contains the protein of interest, which is recruited to the damage ROI. Depending on the filetype of (Movie File), this could be a string (i.e., “EGFP”) or an integer channel ID.
- ROI X Buffer: This input tells the Image Analyzer how many pixels by which to grow or shrink the ROI in the X-dimension. This is used to avoid edge artifacts (i.e., “blurring” of the damage border) from the high intensity laser causing the ROI to be poorly defined, as well as ensuring that the ROI is contained within the nucleus.
- ROI Y Buffer: This input tells the Image Analyzer how many pixels by which to grow or shrink the ROI in the Y-dimension. This is used to avoid edge artifacts (i.e., “blurring” of the damage border) from the high intensity laser causing the ROI to be poorly defined, as well as ensuring that the ROI is contained within the nucleus.
- Pre-irradiation Frames: This is the number of frames from the image stack (Movie File) that are captured prior to DNA damage induction. This information is used to determine the baseline for normalizing the intensity timeseries.
- Additional ROIs: Have the Image Analyzer calculate the kinetics in this number of ROI-sized regions above and below the damage ROI.
- Correct for Drift: Check this box to correct for beam-induced motion of the nucleus.
- Correct for Bleaching: Check this box to apply bleaching correction across the timeseries.
- Submit Job: Click this button to upload the (Movie File) and (ROI File) to the webserver and submit the Image Analyzer task to the queue.

#### *Walkthrough for the Image Analyzer:*

Defining Input Parameters:

1. We begin by providing a name for the workflow, such as “example_job”, to the “Output Title” field.
2. The example imagestack (“example_imagestack.nd2”) and ROI definition files (“example_ROI.tif”) are then uploaded to the “Movie File” and “ROI File” fields.

3. We choose "EGFP" for the tracking channel, as this channel provides the best trace of the nuclear envelope.
4. We also choose "EGFP" for the "Accumulation Channel", as this is the channel of our labeled protein.
5. We apply "-10" and "10" for the "ROI X Buffer" and "ROI Y Buffer", respectively, to cut down on edge-effects of the beam from impacting our total accumulation results.
6. We choose "6" for the value of "Pre-Irradiation Frames", as this is the number of frames captured in the image stack prior to the damage-inducing laser event.
7. We choose 0 additional ROIs, as we are focused on only the damage site.
8. We leave the "Correct for Drift" box checked to ensure that our movies are motion-corrected. It is highly recommended to always correct for drift, unless visual inspection of your image stacks has confirmed that little to no drift occurs over the entirety of the stack.
9. We leave the "Correct for Bleaching" box checked to ensure that our values are not artificially low due to repeated exposures to the excitation laser bleaching our label photocenters.
10. We then click "Submit Job" and wait for the task to complete.\
    - Depending on the size of your uploaded "Movie File", the job may take a moment to initiate while the file is uploaded. During this time, you should not leave the page.
    - Once the upload and submission has completed, you will be brought back to the server's homepage.
    - Progress can be monitored under the "Pipeline Results" tab, but the webserver will send a "Job Completed Successfully" message to your email address upon completion.

<u>Analyzing Output Files</u>

Output files are downloaded from the "Pipeline Results" tab, by clicking the "Download .zip" button. This .zip archive will contain 7 data files, as well as the Image Analyzer output log (here, "example_job.log"). The data files are as follows:

1. "example_imagestack.csv" : The (non-normalized) intensity timeseries for the ROI's and image stack channels that we requested.
2. "example_imagestack_normalized.csv" : The same timeseries information as listed above, except the data has been normalized by the average of the first "Pre-Irradiation Frames" number of frames (here, 6).
3. "example_imagestack_normalized.pdf" : A figure of the normalized intensity timeseries.
    - We use this figure to quickly check for two things:
        - First, we want to make sure that the Image Analyzer accurately generates an accumulation-like curve, which it clearly does in this example (Figure 2).
        - Second, we want to make sure that the accumulation signal-to-noise is sufficient for

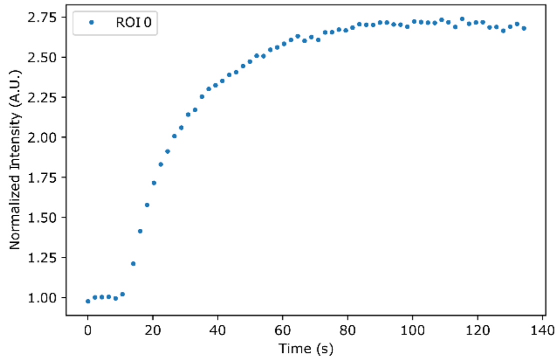


***Figure 2. Normalized accumulation timeseries for the example workflow.*** *The timeseries shows a clear accumulation process, with rapid uptake followed by a plateau.*

fitting diffusion-based models. In this example, signal-to-noise is of no concern (maximum accumulation is ~2.75-fold above baseline, with variations of less than 0.03-fold increase between equilibrium timepoints).

4. "example_imagestack_raw.mp4" : A (non-motion-corrected) movie of the imagestack, with ROI and nuclear envelope overlaid.
5. "example_imagestack_drift_corrected.mp4" : The motion-corrected movie for the image stack, with ROI and nuclear envelope overlaid.
    - Here, we check to make sure that the nuclear envelope nicely traces the visual boundary of the nucleus. In this example, it does indeed trace the nucleus quite well (Figure 3). If this were not the case, then we would have to try using another channel from the image stack for the "Nucleus Tracking Channel". If the nucleus is clearly visible in our movie, but the tracing is still quite poor, then please submit a ticket to: https://github.com/Luger-Lab/Q-FADD/issues

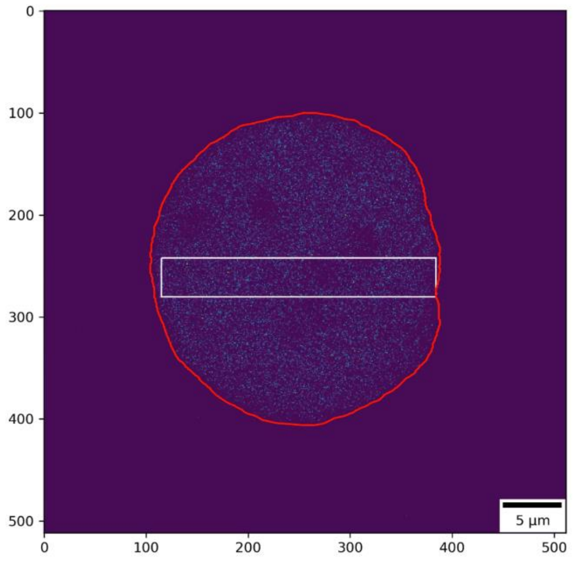


***Figure 3. Nuclear envelope-tracing from the Image Analyzer routine.*** *The red line outlines the nuclear envelope, whereas the white box traces the ROI (damage-induced region). "EGFP" per-pixel intensities reside clearly within the traced envelope.*

6. "example_imagestackNuclMask.txt" : An ASCII file that defines that edges of the nuclear envelope in Cartesian coordinates.
7. "example_imagestackROI.txt" : An ASCII file that defines the rectangular ROI, identifying the DNA damage zone for the diffusion-based modeling routine.

### 2. The Q-FADD Module

After we have confirmed that the nuclear-tracing is appropriate, and that the accumulation curve is of sufficient quality (low noise-to-signal), we can submit the diffusion-based Q-FADD modeling task in one of two ways. First, we could click the "Q-FADD" tab and start a Q-FADD submission (Figure 4). This will require us to upload the Nuclear Mask, ROI Boundary, and Intensity Kinetics files that we just inspected. Alternatively, we could navigate to the "Pipeline Results" tab and select "Start Q-FADD job from output files" for the Image Analyzer task that we just completed. This new Q-FADD task will have an identical "Output Title" as the Image Analyzer task that spawned it.

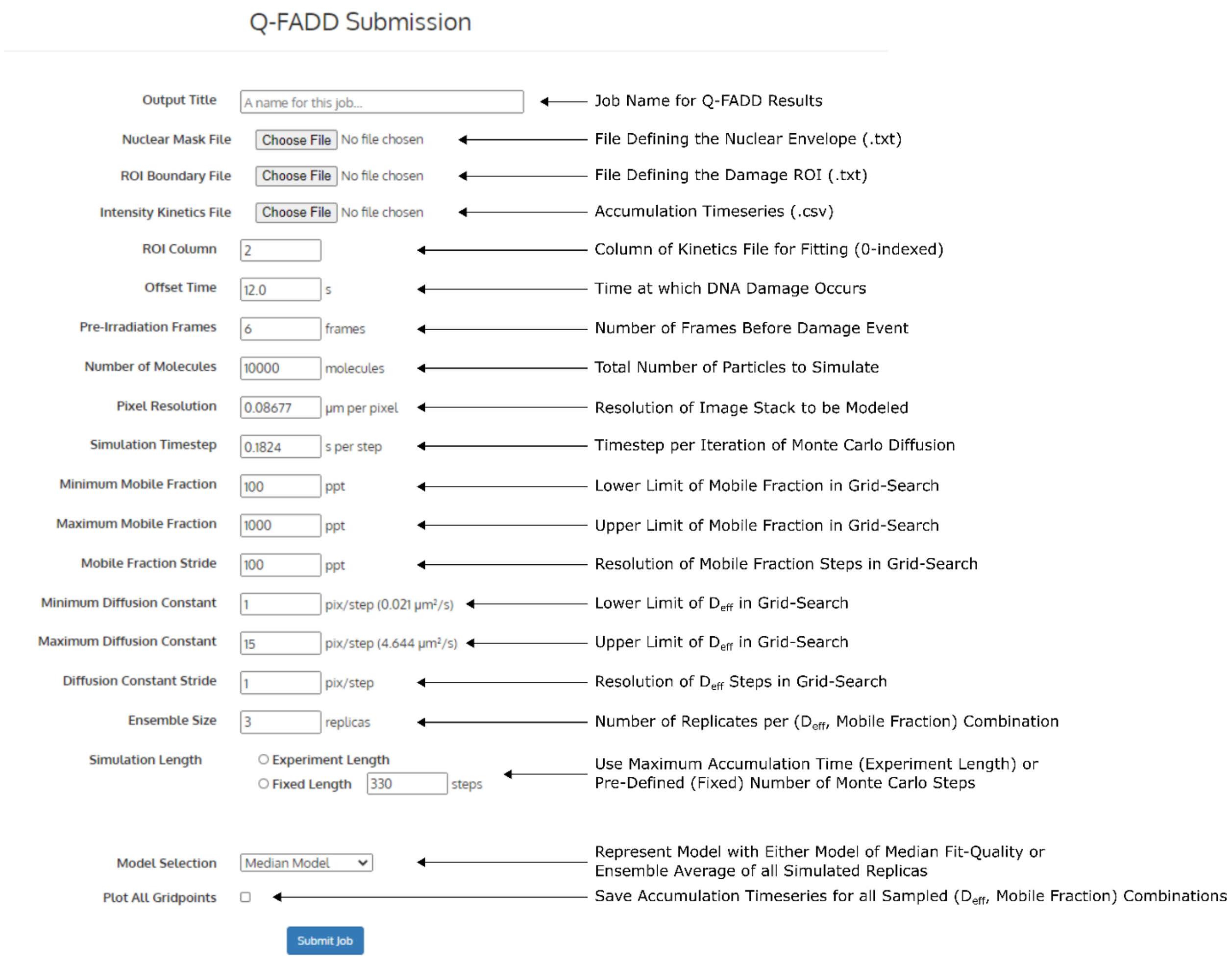


***Figure 4. Snapshot of the Q-FADD submission interface.*** *Nuclear Mask, ROI Boundary, and Intensity Kinetics files can all be uploaded from a user's local computer, or a job can be started from the "Pipeline Results" tab, in which case these fields will be autofilled and require no input from the user.*

Input Parameters:

- <u>Output Title:</u> Name for the output tasks, as will appear in the "Pipeline Results" tab.
- <u>Nuclear Mask File:</u> ASCII file that outlines the nuclear envelope in Cartesian coordinates. (Not required when using the "Start Q-FADD job from output files" option in "Pipeline Results" tab).
- <u>ROI Boundary File:</u> ASCII file that defines the rectangular ROI of the damage site. (Not required when using the "Start Q-FADD job from output files" option in "Pipeline Results" tab).
- <u>Intensity Kinetics File:</u> The .csv file containing the intensity timeseries of interest. (Not required when using the "Start Q-FADD job from output files" option in "Pipeline Results" tab).
- <u>ROI Column:</u> This input identifies which column number (0-indexed) contains the ROI that we would like to fit.
- <u>Offset Time:</u> This input provides the time delay between the t=0 seconds point of the image stack and the t=0 seconds timepoint of the damage event.
- <u>Pre-irradiation Frames:</u> Similar to the Image Analyzer inputs, this tells Q-FADD how many timepoints to use from the "Intensity Kinetics File" to normalize the experimental data.

- Number of Molecules: This is the number of simulated particles to place inside the envelope. This value will be scaled by the "Mobile Fraction" to determine the actual number of diffusing particles in the simulation.
- Pixel Resolution: This tells Q-FADD the pixel resolution (in µm per pixel), with which it scales the effective diffusion constant from pixels per step to $\mu m^2/s$.
- Simulation Timestep: This tells Q-FADD the temporal resolution (in seconds per iteration).
    - This timestep should typically be an order of magnitude higher resolution than your experimental framerate (i.e., 0.2 sec per step for a 2 sec per frame movie) to allow for appreciable sampling of motion between timepoints.
    - However, in the case of proteins that move quite slowly, selecting values more similar to the experimental frame rate may be required to achieve low enough diffusion constant to fit your curve.
        - In these special cases, the number of replicates should also be increased, to counteract the effect of lost temporal sampling.
- Minimum Mobile Fraction: This input determines the lower limit of the "mobile fraction" in the grid search (in parts per thousand – ppt).
- Maximum Mobile Fraction: This input determines the upper limit of the "mobile fraction" in the grid search (in parts per thousand – ppt).
- Mobile Fraction Stride: Using this input, the Q-FADD server will create a distribution of "mobile fraction" values, for the range from minimum to the maximum at every "Mobile Fraction Stride" value.
- Minimum Diffusion Constant: This input determines the lower limit of the effective Diffusion constant in the grid search. This value is defined in pixels per step, and the conversion to $\mu m^2/s$ is also shown.
- Maximum Diffusion Constant: This input determines the upper limit of the effective Diffusion constant in the grid search. This value is defined in pixels per step, and the conversion to $\mu m^2/s$ is also shown.
- Diffusion Constant Stride: This input determines the resolution for the effective Diffusion constant in the grid search.
- Ensemble Size: This parameter defines the number of replicates per ($D_{eff}$, mobile fraction) combination.
    - In our hands, 11 replicates seems to be sufficient for robustly ranking models from all combinations. However, depending on the diffusion rate of the molecule in the cell, as well as signal-to-noise levels for the timeseries, higher values may be required for statistical robustness in separating similarly-performing models.
- Simulation Length: There are two choices when determining how long to run the Monte Carlo simulations of diffusion:
    - Experiment Length: Choosing this option will set the simulated time length to be equivalent to the time required to achieve maximum accumulation.
    - Fixed Length: Alternatively, a fixed number of iterations can be defined. The total simulation length in these cases is then (number of steps) x (Simulation Timestep).
- Model Selection: There are currently two choices for defining model quality for each ($D_{eff}$, mobile fraction) combination:

    - Median Model: The quality-of-fit (and reported simulated accumulation model) is the replicate with the median value for quality-of-fit among the replicates.
    - Average of Models: With this choice, replicate simulations of accumulation for each parameter set are averaged to a single curve, and this average timeseries and fit quality are recorded per grid point (parameter combination).
- Plot All Gridpoints: When selected, the routine will save a PDF figure for all sampled ($D_{eff}$, mobile fraction) combinations in the grid search, allowing users to visually examine fit qualities for all models, and not only the best-identified model.

#### *Walkthrough for the Image Analyzer:*

Defining Input Parameters:

0. Because we used the "Start Q-FADD job from output files" option from the "Pipeline Results" tab, we do not need to define an output title, nor the Nuclear Mask, ROI Boundary, or Intensity Kinetics files, as the webserver will link these automatically.
1. Because we did not define any additional ROIs in the Image Analyzer routine, we know that our intensity values are in the second column of our file (first is the time points). As our column index is 0-indexed, this corresponds to a value of "1" for the "ROI Column" input.
2. We set our "Offset Time" to 12.5 seconds, and the number of pre-irradiation frames to "6" (same value as our Image Analyzer routine).
3. We simulate 10,000 particles in our nuclear envelope.
4. The "Pixel Resolution" of the example_imagestack.nd2 file is 0.08677 µm per pixel, so we use that value here, as well.
5. We set our simulation timestep to 0.2 seconds per step.
6. For the sake of this example, we set our minimum and maximum mobile fractions to 100 and 500 ppt, respectively, and a mobile fraction stride of 100 ppt. This is because we know from previous Q-FADD runs that the best-fit mobile fraction is 300 ppt, so we choose a narrow range here to improve simulation speeds.
    - Users will likely need to sample a wider range of mobile fraction values when modeling their own accumulation data. This limited range was used only because the "answer" was already known.
7. Similarly, we select a narrow range of effective Diffusion constant values (min = 8, max = 13, stride = 1) to improve computational speeds, as we know from previous calculations that the best-fit value is 11 pix/step.
    - Again, users should typically sample a wider range of $D_{eff}$ values.
    - However, wide ranges of $D_{eff}$ and mobile fraction, as well as increased resolution (lower stride values), will become computationally expensive very quickly, if users aren't careful.
        - One strategy to allow for wide searches, but without exceeding computational resources, is to search a wide range of values but at a coarse resolution (strides >100 ppt for mobile fraction, >1 pix/step for Diffusion constant) can identify regions of high quality of fit, and then re-simulating a grid search that spans only these regions at a higher resolution (<100 ppt, and 1 pix/step strides).

8. To further improve computational speeds, we only simulate 3 replicas per grid point. Again, users should expect to simulate at least 11 replicas per parameter combination in order to achieve a statistically robust ranking of sampled models.
9. We elect to use the length of the experiment in order to determine our simulation length, as well as reporting the median model for each parameter set. We did not plot all gridpoints.
10. We then click "Submit Job" and are sent back to the server main page.

Analyzing Output Files:

Output files from the Q-FADD routine are downloaded from the "Pipeline Results" by clicking the "Download .zip" link. In that archive, there are 8 data files, the "example_job.log" file containing the outputs from Q-FADD - including the best-identified parameters for the model of our data ($D_{eff}$ = 11 pix/step, F = 300 ppt, $r^2$ = 0.9876, RMSD = 0.0414) and an "example_job_inputs.par" file, which contains the input parameters for the grid search and Monte Carlo simulations. The 8 data files are as follows:

1. "example_job_all_models.csv" : This .csv file lists sampled $D_{eff}$ (in both pix/step and $\mu m^2/s$) and mobile fraction values, as well as model fitnesses as both $r^2$ and RMSD value (referenced against the experimental curve).
2. "example_job_r2_matrix.dat" and "example_job_r2_matrix.pdf" : ASCII text file and heatmap figure, respectively, for the performance of each sampled grid point in the automated search, using the $r^2$ metric. From the heatmap, users can quickly identify regions of high quality-of-fit (blue and indigo regions, Figure 5A).
3. "example_job_rmsd_matrix.dat" and "example_job_rmsd_matrix.pdf" : ASCII text file and heatmap figure, respectively, for the performance of each sampled grid point in the automated search, using the RMSD metric (referenced against the experimental profile). While qualitatively agreeing with the $r^2$ heatmap, the RMSD-based figure shows higher quantitative resolution for differentiating models of best fit (blue and indigo regions, Figure 5B).

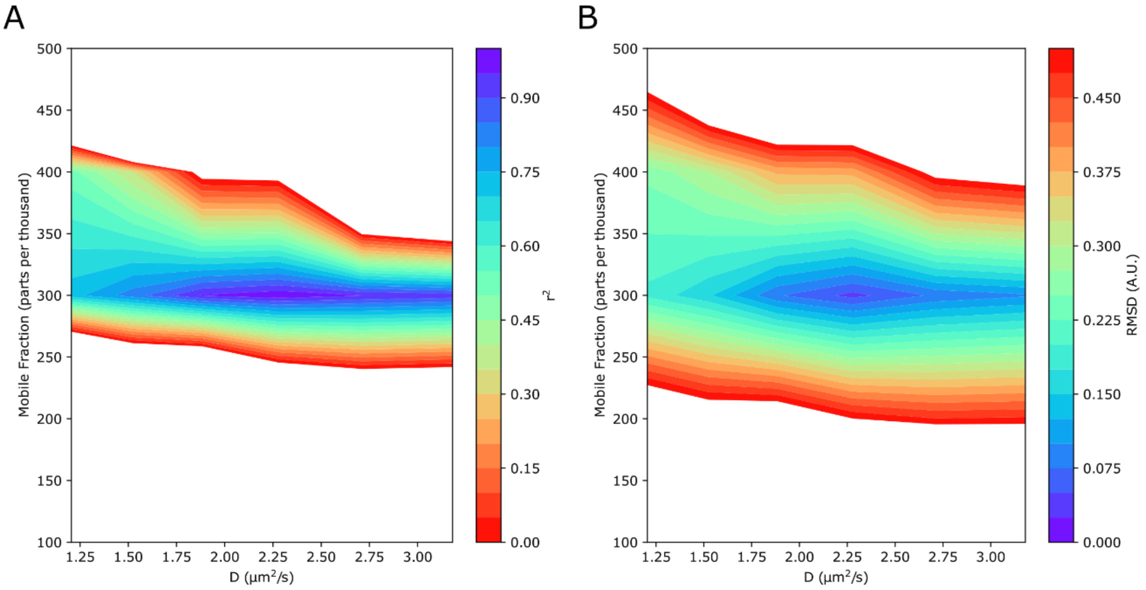


***Figure 5. Heatmaps produced by the example workflow.*** *(A) Heatmap for the automated grid serach, using $r^2$ as the evaluation metric for model quality. (B) Heatmap for the automated grid search, using RMSD (referenced against the experimental timeseries) as the evaluation metric for fitness quality.*

4. “example_job_roi0_intensity_timeseries.csv” and “example_job_Intensity_Timeseries.pdf” : A comma-separated value text file of the model that best fits the experiment and a figure showing the experimental and simulated accumulation profiles (Figure 6, black dots and red line, respectively), respectively. The figure also includes a plot of the fit residuals, giving a means for quickly assessing model performance at different time points.

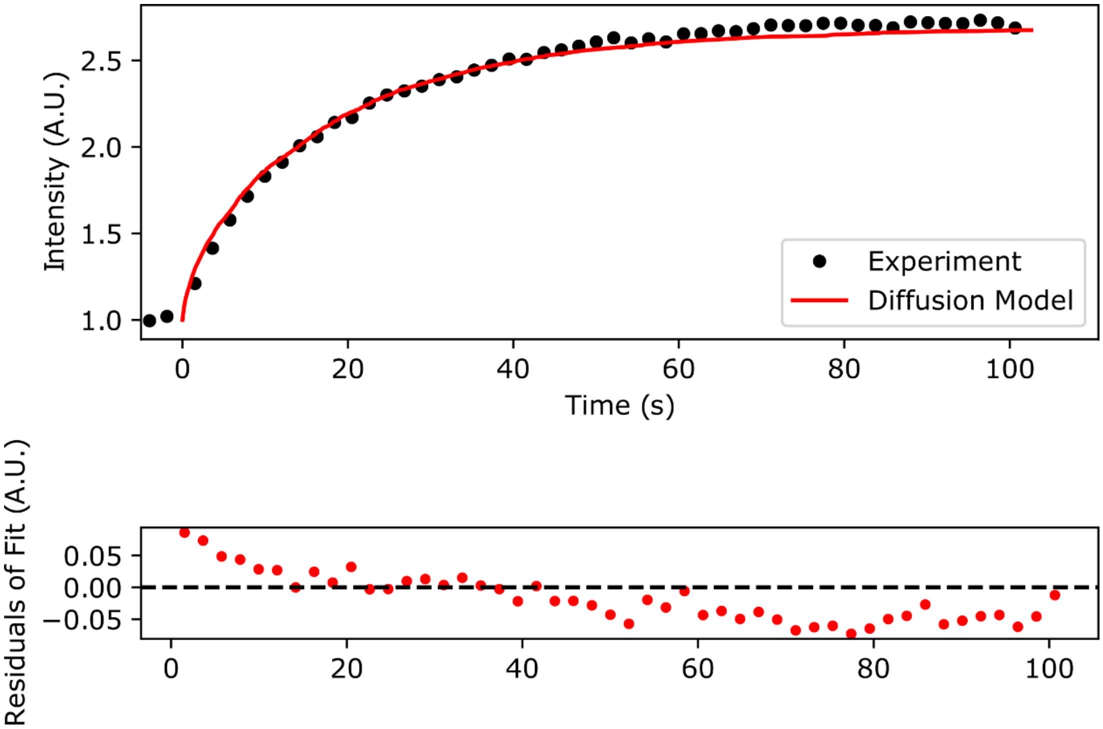


***Figure 6. The “example_job_Intensity_Timeseries.pdf” figure.*** *(top) Comparison between the experimental (black dots) and simulated (red lines) accumulation timeseries. Note, that the timepoints have been translated such that t=0 sec represents the time at which DNA damage was introduced. (bottom) Residuals of the fit between the experimental and simulated accumulation profiles.*

5. “example_job_Model_Nucleus_with_ROI.pdf” : A figure showing the overlay between the nuclear envelope and the damage ROI, as drawn according to the “Nuclear Mask” and “ROI” text files.
