## Supplementary material for "Automated Modeling of Protein Accumulation at DNA Damage Sites using qFADD.py": example_files.zip: single_file_example.bash

```
#!/bin/bash

#The commands below will run the full qFADD.py pipeline on a single file, including both imagestack analysis and diffusion-based modeling

#This is the imagestack analysis command. Will take several minutes to run.
image_analyzer.py -m_file example_imagestack.nd2 -roi_file example_ROI.tif -o example_job/ -nchan 'EGFP' -pchan 'EGFP' -x_buff "-10" -y_buff "10" -n_rois 0 -pre_frames 6 --drift_correct --bg_correct

#We then enter the "example_job" folder from the image_analyzer.py output and then run qFADD.py on the files in that folder
cd example_job/

#This will run qFADD.py on 20 processors (-np 20), so you should adjust the number of processors to match the capabilities of your own local environment.
mpirun -np 20 qFADD.py \
    -mf ./example_imagestackNuclMask.txt \  #This is the ASCII file that defines the nuclear envelope
    -roif ./example_imagestackROI.txt \     #This is the ASCII file that defines the rectangular ROI
    -kf ./example_imagestack.csv \          #This is the .csv file that gives accumulation intensity vs time
    -roic 1 \                               #This tells qFADD which column of the above .csv contains the timeseries that we are fitting
    -out  example_imagestack_qFADD \        #This is the prefix for the collection of output files.
    -t0   12.5 \                            #This tells qFADD the time at which the damage-inducing laser was engaged (defines the t=0 point for the accumulation process)
    -norm 6 \                               #This tells qFADD how many frames to average for the normalization factor. Should be identical to the number of pre-irradiated frames used in the image_analyzer.py function.
    -nmol 10000 \                           #This is how many labeled-proteins to simulate. On the order of 10^4 seems sufficient in our tests, so far.
    -mobile_min 100 \                       #This sets the lower bound for the "mobile fraction" dimension of the grid search.
    -mobile_max 1000 \                      #This sets the upper bound for the "mobile fraction" dimension of the grid search.
    -mobile_step 100 \                      #This sets the stride for the "mobile fraction" dimension, will generate grid points every "mobile_step" between "mobile_min" and "mobile_max".
    -pix 0.08677 \                          #This is the um per pixel resolution of the experimental imagestack (can be found from output of image_analyzer.py, if unknown).
    -Dmin 1 \                               #This is the lower limit for the "effective Diffusion coefficient" dimension of the grid search. (in pix/step)
    -Dmax 15 \                              #This is the upper limit for the "effective Diffusion coefficient" dimension of the grid search. (in pix/step)
    -Dstep 1 \                              #This is the stride for the "effective Diffusion coefficient" dimension (in pix/step), will generate grid points every "Dstep" between "Dmin" and "Dmax".
    -ensemble_size 11 \                     #Sets the number of replicates per grid point, to establish statistical robustness. 11 replicates have been found to work fairly consistently, in our hands.
    --find_plat \                           #Set the total number of steps for the simulation to match the time to achieve maximum total accumulation in the experimental timeseries.
    -tstep 0.2 \                            #This sets the timestep of the diffusion models (in sec per iteration).
```
