## Supplementary figures and images for "Automated Modeling of Protein Accumulation at DNA Damage Sites using qFADD.py"

### example_imagestack_normalized.pdf

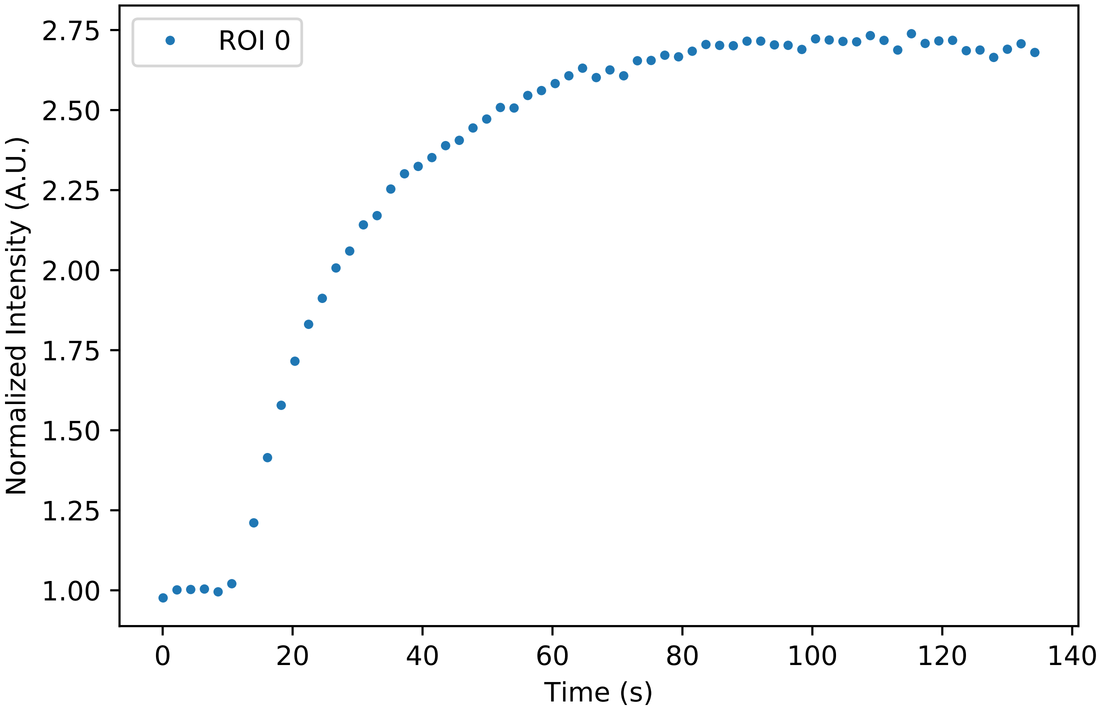

### example_imagestack_normalized.png

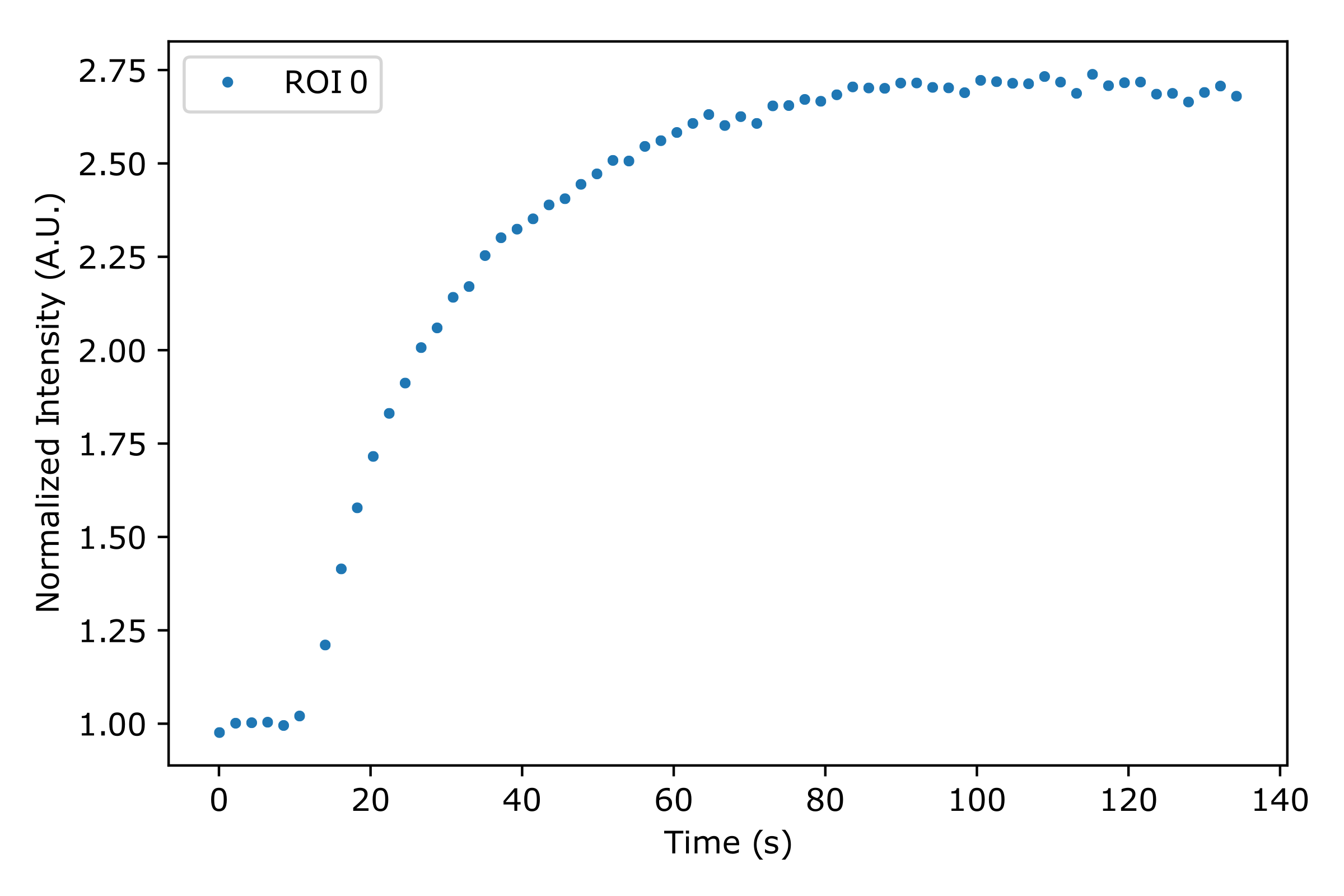

### example_imagestack_qFADD_Intensity_Timeseries.pdf

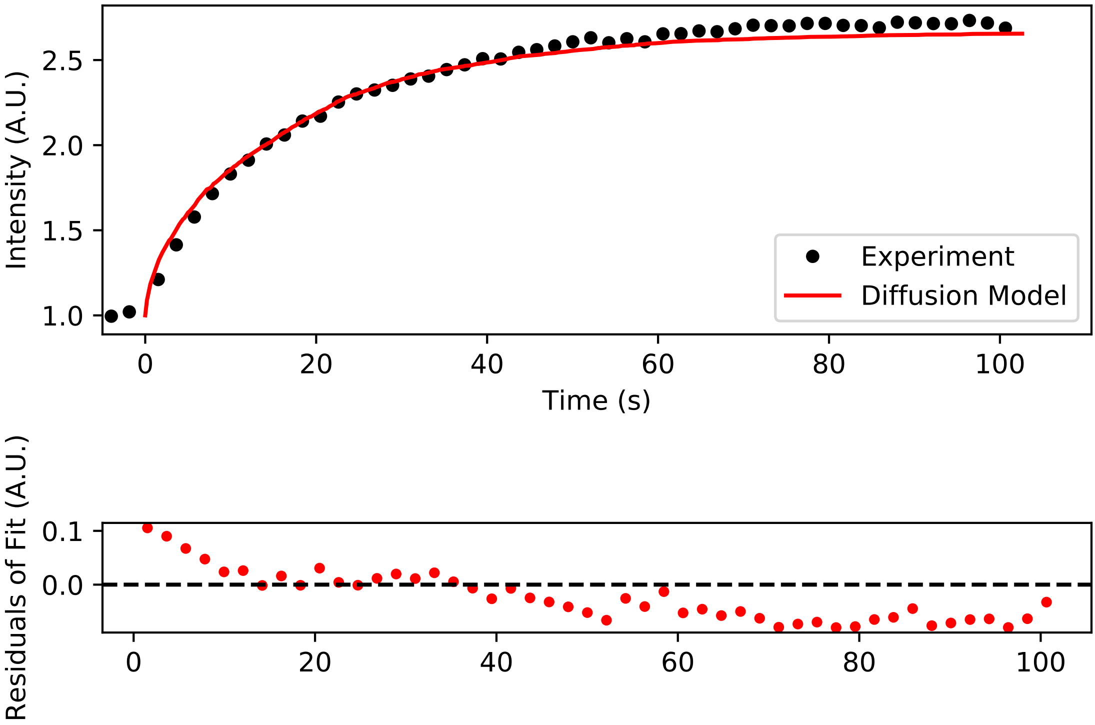

### example_imagestack_qFADD_Model_Nucleus_with_ROI.pdf

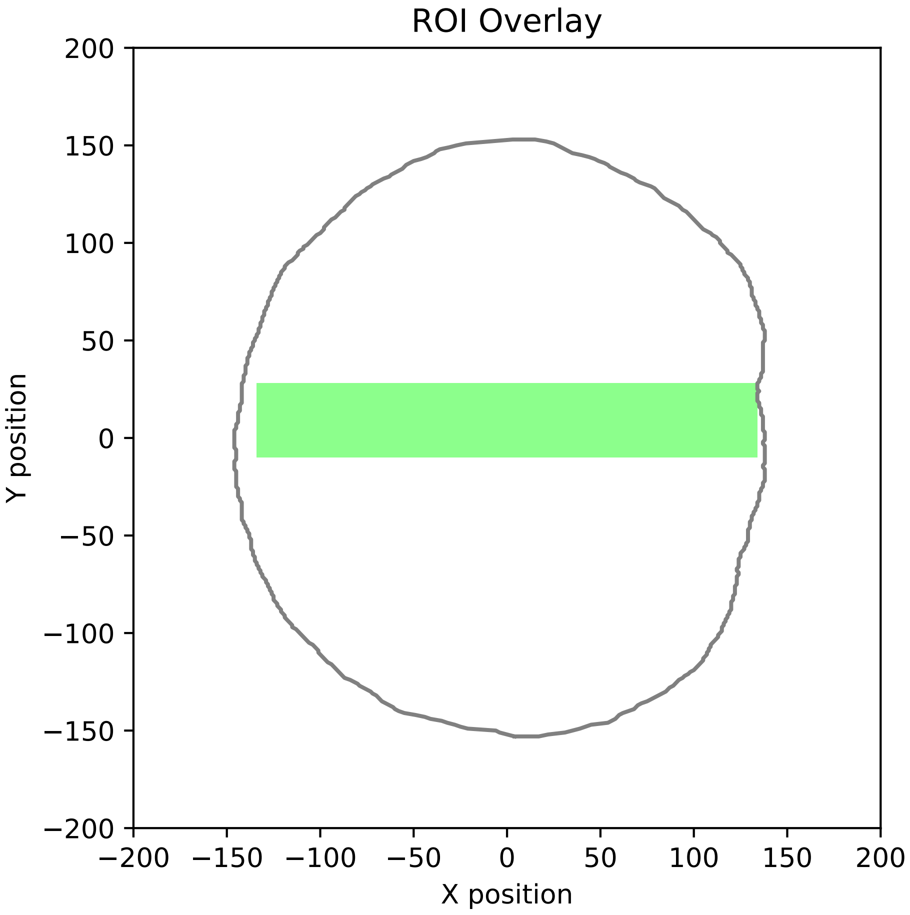

### example_imagestack_qFADD_r2_matrix.pdf

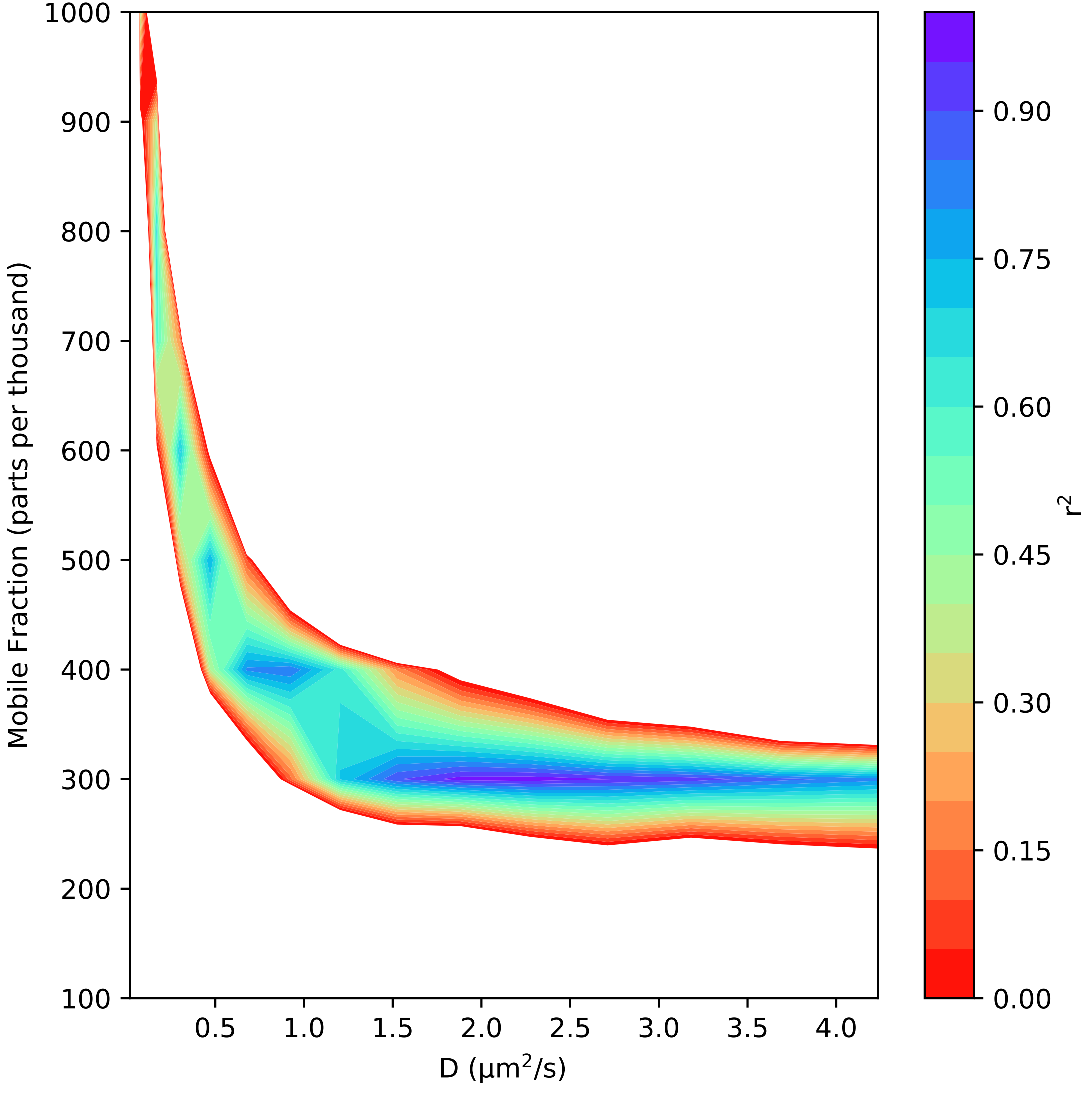

### example_imagestack_qFADD_rmsd_matrix.pdf

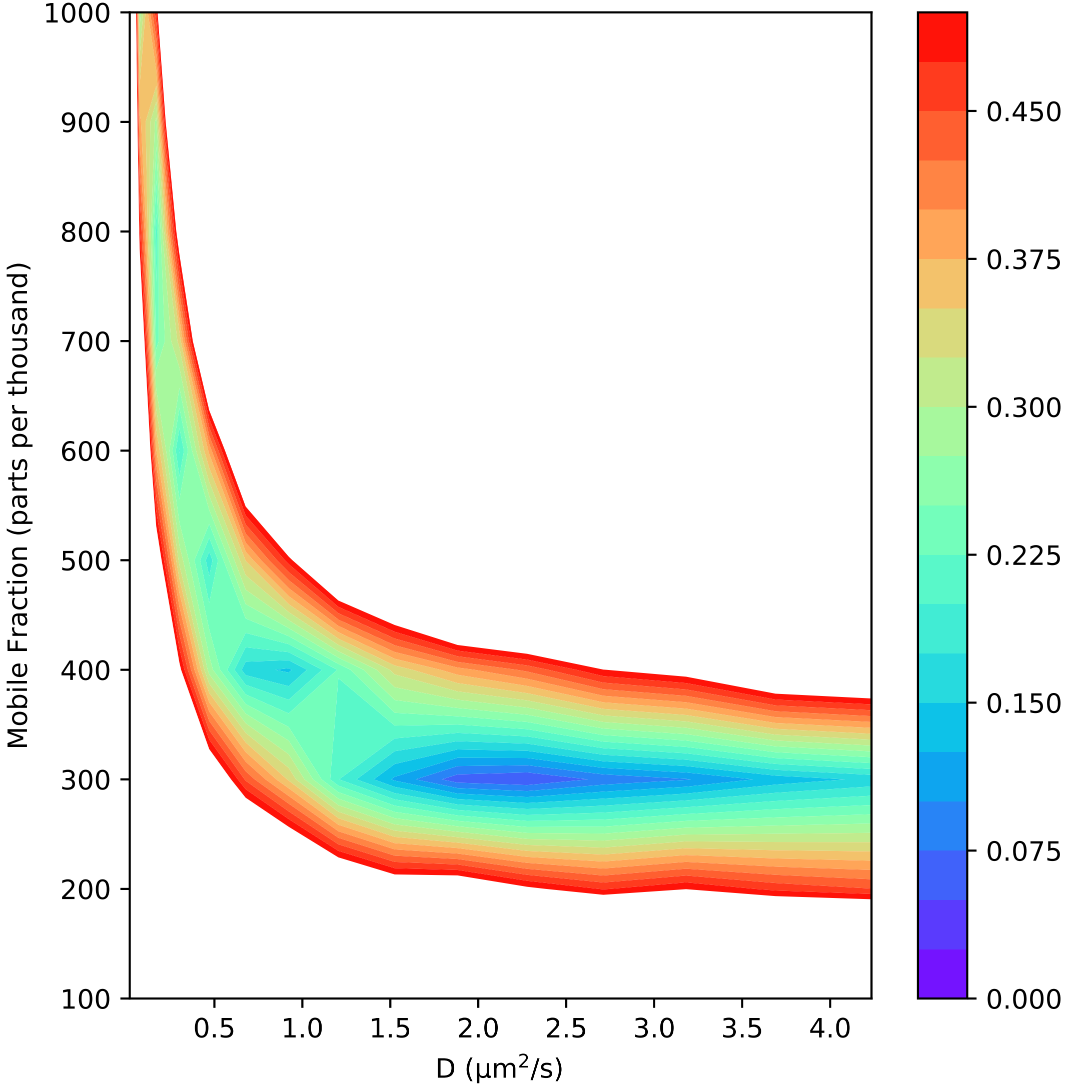

### example_ROI.tif

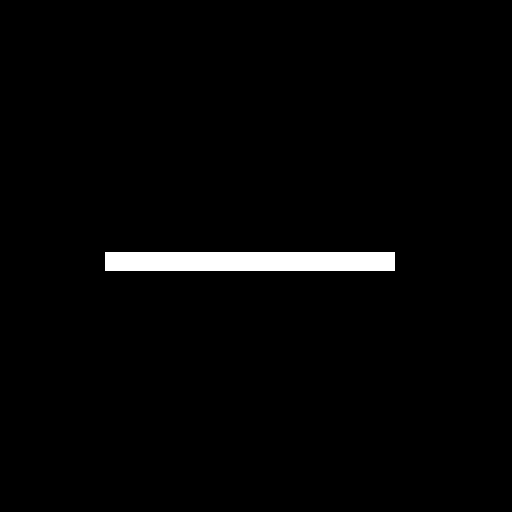
